# Continuous partitioning of neuronal variability

**DOI:** 10.1101/2025.07.23.666404

**Authors:** Anuththara Rupasinghe, Adam S. Charles, Jonathan W. Pillow

## Abstract

Neurons exhibit substantial trial-to-trial variability in response to repeated stimuli, posing a major challenge for understanding the information content of neural spike trains. In the visual cortex, responses show greater-than-Poisson variability, whose origins and structure remain unclear. To address this puzzle, we introduce a continuous, doubly stochastic model of spike train variability that partitions neural responses into a smooth stimulus-driven component and a time-varying stochastic gain process. We applied this model to spike trains from four visual areas (LGN, V1, V2, and MT) and found that the gain process is well described by an exponentiated power law, with increasing amplitude and slower decay at higher levels of the visual hierarchy. The model also provides analytical expressions for the Fano factor of binned spike counts as a function of timescale, linking observed variability to underlying modulatory dynamics. Together, these results establish a principled framework for characterizing neural variability across cortical processing stages.

## 1 Introduction

Neural responses exhibit trial-to-trial variability, even in response to repeated presentations of the same stimulus [1–5]. Several hypotheses have been proposed for the source of this variability, including noise in synaptic transmission [6], fluctuations in balanced excitation and inhibition [7], noisy peripheral sensors [8], recurrent dynamics [9], neural adaptation [10] and synaptic plasticity [11]. To assess the impacts of variability on neural coding, researchers have developed a variety of statistical models that seek to capture both the stimulus dependence and stochasticity of neural spike responses [12–17]. The simplest such model for spike train data is the Poisson process [18, 19], which assumes that spike count variability in disjoint time intervals is independent. Under this model, the variance of the binned spike counts is equal to the mean for any choice of time bin. However, a large literature has shown that this relationship does not hold precisely in many brain areas. Rather, spike count variance commonly exceeds the mean, a phenomenon known as “overdispersion” [1, 7, 20–30].

To account for super-Poisson response variability in the visual pathway, Goris et al. [1] introduced the “modulated Poisson” model, which partitions neural variability into a stimulus-driven component and a modulatory gain component. Specifically, the model assumes that spike counts are Poisson distributed, with a mean determined by the product of two terms: a stimulus-related component that reflects the neuron’s stimulus tuning, and a stimulus-independent stochastic gain that varies randomly across trials (Figure 1A). Goris et al. [1] showed that this model provides an accurate account of spike count variability across the early visual pathway, with the degree of overdispersion linked to the strength of the modulatory component.

**Figure 1.**
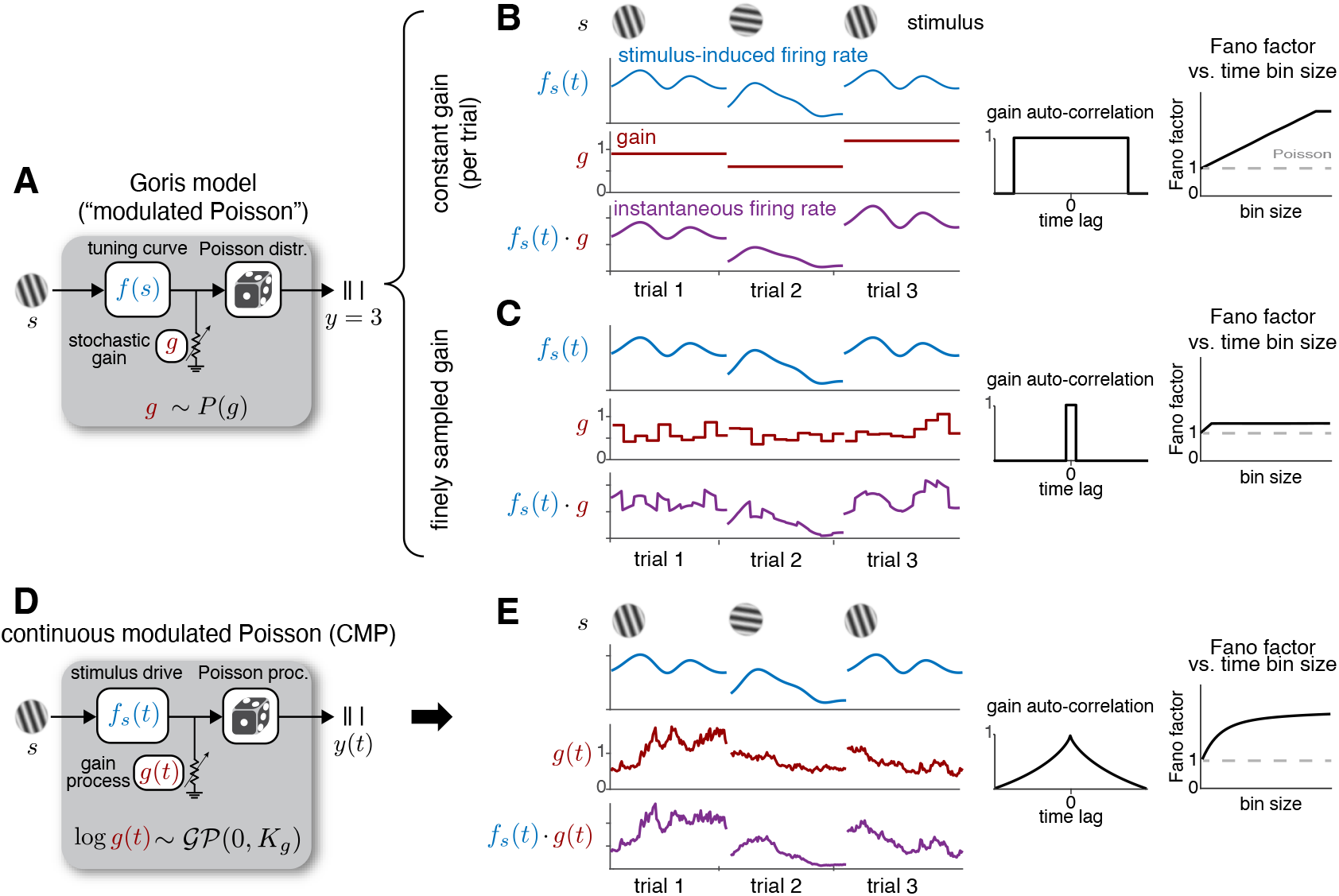
(A) Schematic of the Goris modulated Poisson model [1], in which spike counts (*y*) arise from a Poisson process with rate given by the product of two components: a stimulus-dependent drive *f* (*s*) and a stimulus-independent gain *g*. The gain *g* is a scalar and is independently drawn across trials. (B, C) Two continuous-time extensions of the Goris model. (B) Constant gain model: assumes that the sampling interval of the gain equals the trial duration. Left: stimulus-induced rate, gain process, and instantaneous firing rate for three example trials. Middle: gain autocorrelation function. Right: Fano factor as a function of bin size (dashed line indicates the baseline Poisson model). (C) Independent gain model: assumes that the sampling interval of the gain is much shorter than the length of a single trial. Panels are organized as in (B). (D) Schematic of the proposed Continuous Modulated Poisson (CMP) model. Here, spike trains *y*(*t*) are generated by a Poisson process whose rate is the product of a time-varying stimulus drive *f*_*s*_(*t*) and a continuous-time stochastic gain process *g*(*t*). The gain process *g*(*t*) follows a Gaussian process (GP) with temporal correlations governed by an exponentiated power law kernel, and is independently drawn across trials. (E) Panels are organized as in (B), for the CMP model.

However, in the original modulated Poisson formulation [1], the modulatory gain is represented as a scalar random variable associated with a counting window or trial. Thus, while the model provides an effective account of spike count overdispersion, it does not explicitly model gain as a continuously timevarying process within a trial. This limits its ability to characterize how gain fluctuations evolve across subtrial timescales. Related work has further examined the dependence of overdispersion on counting-window duration and related time-dependent extensions of the modulated Poisson framework [31, 32]. Here, we build on this line of work by introducing an explicit continuous-time model of within-trial gain fluctuations.

To clarify this distinction, we start by considering two possible continuous-time interpretations of the modulated Poisson framework of Goris et al. [1]. In both cases, we allow the stimulus component to be a stimulus-dependent, time-varying Poisson firing rate and impose a smoothness prior on this stimulus drive, so that the comparison focuses on different assumptions about the temporal structure of the gain. This smoothness prior distinguishes our implementation from related time-dependent variants of the modulated Poisson framework [31, 32], and provides a regularized estimate of the time-varying stimulus component. We then consider two different assumptions about the temporal sampling of the stochastic gain variable. First, the gain may be constant across the full stimulus presentation or trial (Figure 1B). In this case, the gain is independently drawn for each trial, producing a gain process with broad temporal correlations and a Fano factor (the spike-count variance divided by the mean spike count) that grows approximately linearly with the size of the counting window (Figure 1B, right). As a second alternative, the gain may be resampled independently across time bins on a timescale much shorter than the length of a single trial (Figure 1C). In this case, the gain autocorrelation is narrow, and the Fano factor curve saturates for bin sizes larger than the gain sampling interval (Figure 1C, right).

These continuous-time interpretations of the Goris model rely on the assumption that the modulatory gain is constant during each sample interval and then jumps discontinuously between intervals. Such piecewise-constant gain dynamics are unlikely to capture the smooth temporal structure of modulatory fluctuations in biological systems. Moreover, these models have limited ability to capture complex relationships between time bin size and Fano factor, given that they primarily produce curves that grow linearly and then saturate (Figure 1B,C).

To overcome these limitations, we introduce the continuous modulated Poisson (CMP) model, a flexible extension of the Goris model to continuous-time spike responses. The CMP model describes spike trains as arising from a Poisson process with a rate given by the product of two continuous-time processes: (1) a “stimulus-driven” component, which describes the firing rate fluctuations induced by the stimulus; and (2) a modulatory “gain” component, which captures stimulus-independent trial-to-trial variability using a stochastic process (Figure 1D). Both components are modeled as exponentiated Gaussian processes (GPs), providing a flexible framework for capturing smooth, continuous dynamics with tunable temporal structure [33–35].

To more precisely capture the temporal correlations in the modulatory gain process, we introduce a novel covariance function that allows these correlations to decay more slowly than those described by the standard radial basis (or “Gaussian”) kernel. We refer to this as the Exponentiated Power Law (EPL) kernel, which models autocorrelation decay according to an exponentiated power law with exponent *q* ≤ 2 (Figure 1E, middle). This added flexibility enables the model to more accurately capture nonlinear relationships between bin size and Fano factor (Figure 1E, right) compared to previous approaches. Moreover, the CMP framework provides closed-form expressions for how the Fano factor scales with time bin size, offering a direct way to probe how variability accumulates and transforms across different temporal resolutions.

We applied the CMP model to spike train recordings from four visual areas (LGN, V1, V2, and MT) [1, 36] and uncovered novel insights into variability across the visual hierarchy. In particular, we observed a systematic increase in the magnitude of gain fluctuations with progression along the visual hierarchy, consistent with prior observations [1]. Furthermore, comparisons of the inferred EPL kernels across visual areas revealed that gain autocorrelation exhibits slower temporal decay at higher levels of the visual hierarchy. A comparison of the variance of the stimulus drive with the variance of the modulatory gain process across neurons further revealed that these two components tend to be negatively correlated in all visual areas. These findings support existing hypotheses while providing new insight into variability structure and demonstrate that the CMP model outperforms existing methods in capturing time-resolved overdispersion.

In summary, this study makes five main contributions. (1) We extend classical variability partitioning approaches from spike-count models to a continuous-time spike-train framework, allowing neural variability to be characterized as a function of temporal scale rather than fixed counting windows. (2) We introduce the Exponentiated Power Law (EPL) kernel, which models slow, heavy-tailed correlations in modulatory gain and links them to distinct timescales of variability. (3) We derive analytic expressions for the Fano factor as a function of time bin size, enabling variability to be partitioned across temporal scales within a single unified model. (4) We develop a scalable inference framework that jointly estimates stimulus-driven and modulatory components directly from spike train data. (5) By applying CMP to large-scale recordings across multiple stages of the visual hierarchy, we reveal systematic trends in gain dynamics and stimulus modulation, providing new insights into how variability is structured and propagated across cortical processing stages. Altogether, CMP preserves the interpretability of classical variability-partitioning models while extending them to continuous-time spike-train data, enabling time-resolved analysis of gain dynamics across stimulus conditions and brain areas.

## 2 Results

### Continuous modulated Poisson (CMP) model

Our proposed model describes spike trains using a doubly stochastic point process model [37], in which the firing rate of a Poisson process is itself a stochastic process. Specifically, we model a spike train *y*(*t*) elicited in response to a stimulus *s* using a Poisson process:

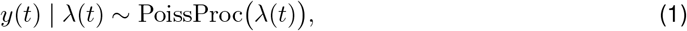

where the firing rate, or “conditional intensity”, *λ*(*t*), is a random variable defined as the product of two terms:

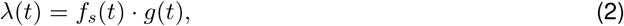

with *f*_*s*_(*t*) representing the stimulus-dependent component that captures stimulus-locked fluctuations in firing rate, and *g*(*t*) denoting a stochastic gain process that accounts for stimulus independent fluctuations in neuronal excitability (Figure 1D-E).

We assume that the stimulus component *f*_*s*_(*t*) varies independently across stimuli and evolves smoothly in time. To express this smoothness, we place a Gaussian process (GP) prior over the log firing rate:

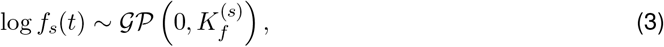

where 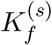 is the standard radial basis function (RBF) covariance function, which describes the covariance of fluctuations in log firing rate at arbitrary points in time [33]. Specifically, the covariance between the log firing rate at any two time points *t*_1_ and *t*_2_ during stimulus presentation interval [0, *T*] is given by:

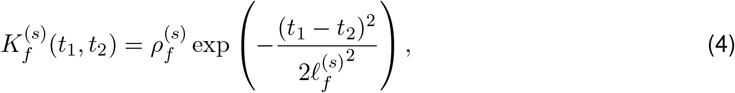

where 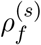 and 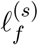 are hyperparameters controlling, respectively, the variance and temporal smoothness (length scale) of fluctuations in the log firing rate. To flexibly capture diverse tuning profiles, each stimulus condition is assigned its own set of stimulus-related hyperparameters.

The gain process *g*(*t*) is modeled analogously as an exponentiated Gaussian process:

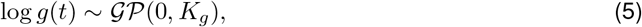

where *K*_*g*_ is the covariance function that characterizes temporal correlations in the gain process. Unlike the stimulus component, the gain is a latent variable that varies independently across repeated presentations of the same stimulus, thereby inducing greater-than-Poisson variability in the spike counts. For a fixed stimulus *s* = *s*^∗^ presented *N* times, the model includes a shared stimulus component *f*_*s*_∗ (*t*) and an independent gain process *g*_*i*_(*t*) for each repeat *i* ∈ {1, …, *N*}. Also, in contrast to the stimulus drive, because the gain process reflects stimulus-independent variability, its hyperparameters are shared across all stimulus conditions. This formulation provides a unified and parsimonious framework for modeling trial-to-trial variability across stimuli within a single generative model.

To more accurately model the stochastic structure of real neural spike trains, we introduce a novel covariance function for the gain process, referred to as the Exponentiated Power Law (EPL) kernel:

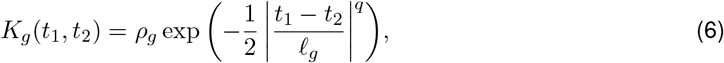

where *ρ*_*g*_ denotes the marginal variance, *ℓ*_*g*_ is the temporal length scale, and *q* is the power-law exponent controlling the rate of correlation decay over long timescales. We show in the Appendix 1 that this kernel defines a valid covariance function for 0 *< q* ≤ 2. Because the gain reflects stimulus-independent variability, its hyperparameters are shared across all stimulus conditions. This formulation provides a unified and parsimonious generative model that captures both smooth, stimulus-locked activity and structured trial-to-trial variability across stimuli.

Figure 2 illustrates example gain covariance functions under different settings of the hyperparameters *ℓ*_*g*_ (length scale) and *q* (power-law exponent). When *q* = 2 (right column), the EPL kernel reduces to the standard RBF covariance function defined in Equation 4. For *q <* 2, the correlations exhibit heavier tails than in the RBF case, decaying more slowly at long time lags. In the limiting case of *ℓ*_*g*_ → ∞ or *q* → 0, the gain becomes effectively constant over each stimulus presentation interval [0, *T*], corresponding to the continuous-time Goris model shown in Figure 1B.

**Figure 2.**
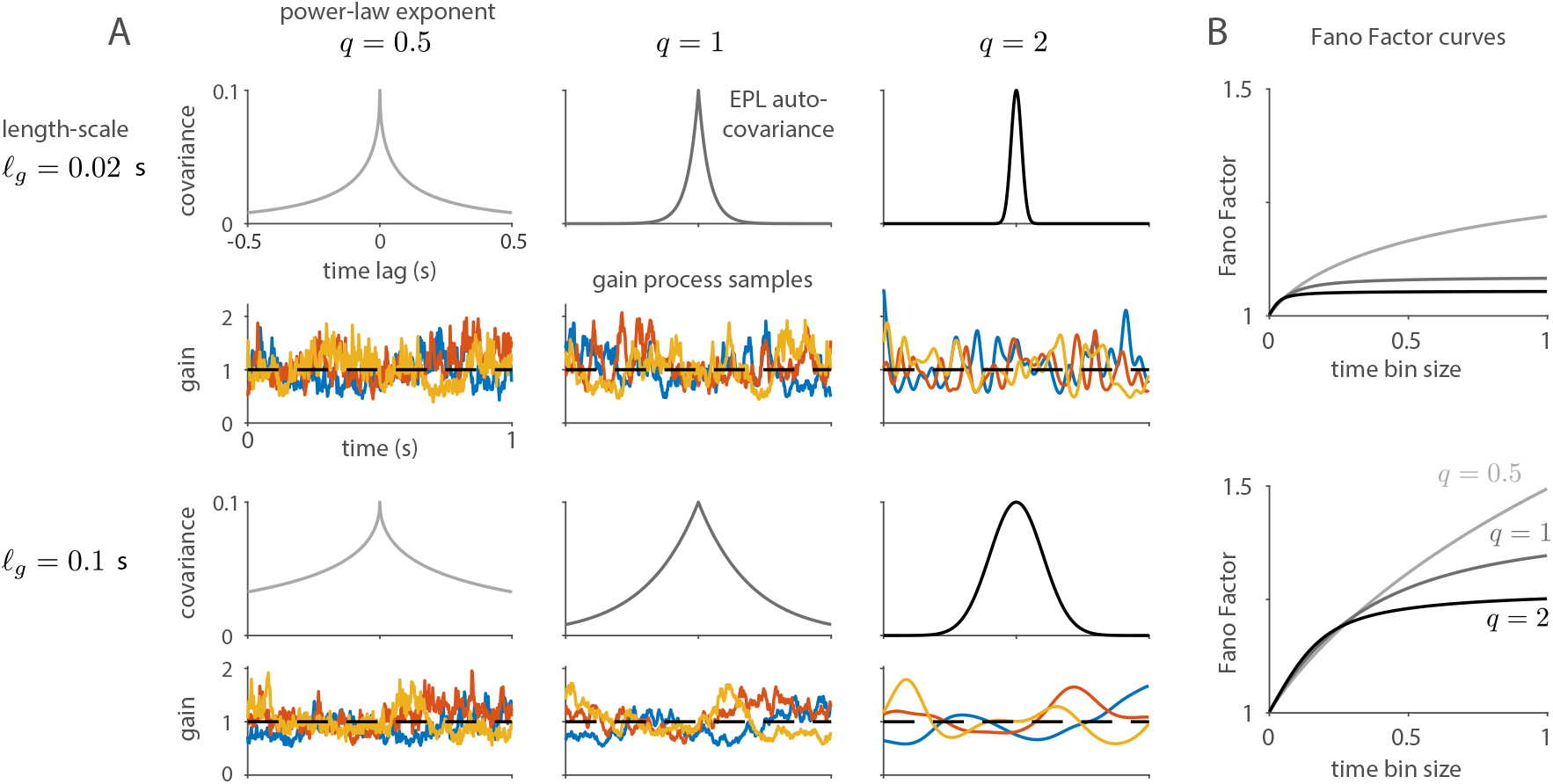
Illustration of different settings of the proposed Exponentiated Power Law (EPL) covariance function with marginal variance *ρ*_*g*_ = 0.1. **(A)** EPL auto-covariance functions with different settings of length-scale (top row: *ℓ*_*g*_ = 0.02, bottom row: *ℓ*_*g*_ = 0.1) and power-law exponent (columns left to right: *q* = 0.5, *q* = 1, *q* = 2). Colored traces below each auto-covariance show three sample gain processes *g*(*t*), obtained by sampling from the corresponding GP and exponentiating. **(B)** Fano Factor curves showing how Fano Factor changes as a function of the bin size used to count spikes, for EPL covariances in the top and bottom rows in (A), respectively.

### Partitioning variability with CMP

Under the CMP model, spike train variability can be partitioned into a Poisson component, which is equal to the mean, and a modulatory component, which controls the degree of overdispersion. The continuous-time formulation of the CMP model allows us to compute this partitioning for arbitrary time intervals, and thus to analytically characterize how overdispersion grows with time bin size. In the case of a non-fluctuating stimulus component 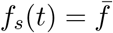, the mean spike count in a time bin of size Δ is given by:

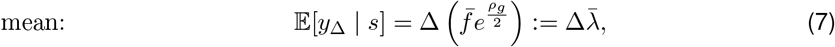

where 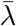 denotes the mean firing rate under the model. The spike count variance is then given by:

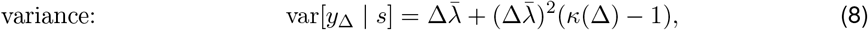

where

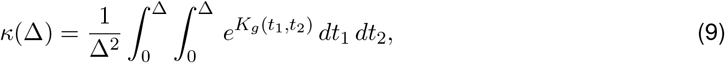

is the average of the exponentiated covariance function exp(*K*_*g*_) over all pairs of times in the interval [0, Δ]. (Note: this formula holds for any covariance function, not just the EPL covariance function we use for *K*_*g*_). From these two terms, the Fano factor as a function of bin size can be computed as the ratio of variance to mean:

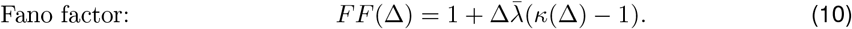

This shows that the Fano Factor approaches 1 in the limit of small bin size Δ, low firing rate 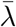, or when *κ*(Δ) approaches 1, which occurs when the variance of the gain process *ρ*_*g*_ goes to zero. Note that *κ*(Δ) is a decreasing function of Δ, reflecting the decay of correlations in the gain process, and it remains constant only for a gain process with infinite lengthscale (for example, Figure 1B). Thus, the Fano factor of the CMP model grows at most linearly with time bin size. Corresponding expressions for the general case, where *f*_*s*_(*t*) is a fluctuating stimulus component, are provided in the Methods and Materials (see Appendix 2 for derivations).

### Model fitting

To fit the CMP model to data, we need to compute the probability over spike counts in finite-sized time bins, given that spike trains are generally measured at a finite sampling rate. Computing this probability involves integrating a conditionally Poisson spike count probability over all realizations of the gain process *g*(*t*):

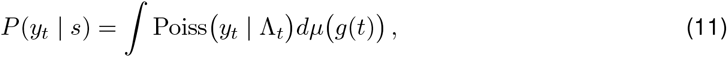

where 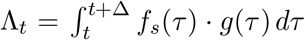 is the expected spike count in the bin [*t, t* + Δ] given the stimulus *s* and gain process *g*(*t*), and *µ(g*(*t*)) is the probabilistic measure of the gain process under the Gaussian process prior (Equation 5). However, this (infinite-dimensional) integral cannot be computed in closed form, since it involves the product of Poisson and Gaussian distributions. Thus, we introduce an efficient variational inference procedure [38], which allows us to optimize the hyperparameters governing the GP covariance functions and infer the stimulus components *f*_*s*_(*t*) for each stimulus (see Methods and Materials). The fitting method also provides a log-Gaussian approximation to the posterior over the gain process *g*(*t*) for each trial, allowing us to visualize the most likely modulatory stochastic gain trajectory for each repeat of a given stimulus.

To validate our inference procedure, we applied it to a synthetic dataset generated from the CMP model, as shown in Figure 3. We simulated spiking activity over a duration of *T* = 5 s for *S* = 3 stimulus conditions, with *N* = 50 repeated trials per stimulus, following the CMP generative model. We then fit the CMP model to the simulated spike trains using a temporal grid size of *dt* = 0.05 s and estimated both the latent processes and GP hyperparameters (see Methods and Materials for details). Figure 3A illustrates the true and inferred stimulus-drive and gain processes for two representative trials from two different stimulus conditions, along with the corresponding firing rates and observed spike counts. Figure 3B compares the true and inferred Gaussian process hyperparameters. These results show that the proposed inference framework accurately recovers the ground-truth stimulus drives, gain processes, and GP hyperparameters on a discretized time grid. Note that the model itself is defined in continuous time and can therefore be consistently applied to data measured at arbitrary temporal resolutions.

**Figure 3.**
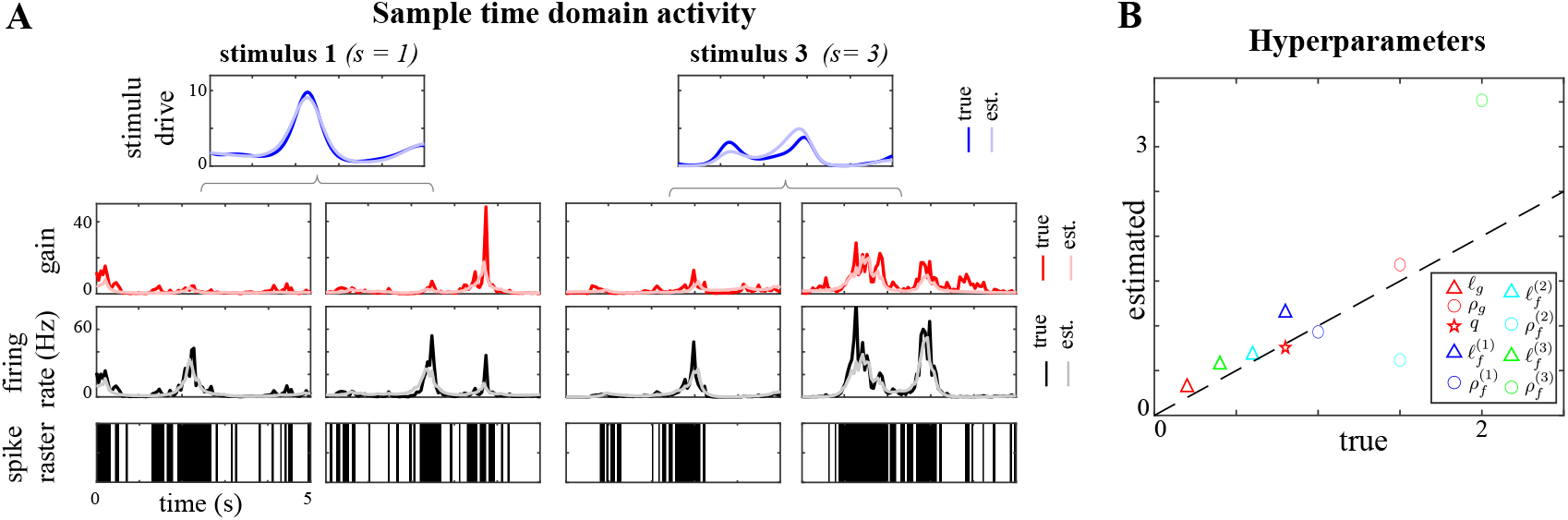
Simulation-based validation of CMP inference. **(A)** From top to bottom: true and inferred stimulus-driven components for stimulus 1 and stimulus 3; true and inferred gain processes for two representative trials from each stimulus condition; corresponding true and inferred firing rates; and observed spike counts. **(B)** Comparison of true and inferred gain hyperparameters *ℓ*_*g*_, *ρ*_*g*_, *q* and stimulus-drive hyperparameters 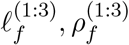.

### Continuous partitioning of variability in the macaque V1

We applied our method to partition response variability in five populations of neurons recorded from the superficial layers of the macaque primary visual cortex (data from [36]). These datasets were collected from three monkeys using drifting grating stimuli presented at *S* = 72 equally spaced orientations. Each grating was displayed for 1280 ms, followed by a 1280 ms blank period, resulting in a total trial duration of 2.56 s. Each grating orientation was repeated *N* = 50 times. In total, we analyzed spiking activity from 654 isolated units (147, 113, 113, 133, and 148 neurons across the five datasets).

Figure 4A shows the inferred time-domain activity of a representative neuron from dataset 1, using a temporal grid resolution of *dt* = 0.02 s, for three different drift orientations (0^0^, 105^0^, and 280^0^). The rows, from top to bottom, show the PSTH derived from the data, the estimated stimulus drive, and, for two different trials per orientation, the inferred gain processes, firing rates, and observed spike rasters, followed by the cross-trial gain mean and cross-trial gain variance. While the estimated stimulus drive closely matches the PSTH, the inferred gain processes capture trial-specific temporal fluctuations that are clearly visible in the spike rasters. Notably, when the neuron is strongly driven by the stimulus (middle column), the instantaneous per-trial firing rate is dominated by the stimulus drive. In contrast, when the neuron is weakly driven (left column), the stochastic gain process plays a larger role in shaping the per-trial firing rate. Importantly, the cross-trial gain mean is relatively flat, and does not resemble the inferred stimulus drive or exhibit clear stimulus-locked temporal structure. This suggests that trial-specific gain fluctuations largely average out across repeats, consistent with the interpretation that the gain process captures random trial-to-trial variability rather than stimulus-driven activity. In contrast, the cross-trial gain variance shows a general reduction during the stimulus presentation compared to the inter-trial interval, indicating that gain variability is quenched during stimulus presentation. These observations indicate that the CMP model meaningfully partitions response variability across stimulus conditions. Figure 4B shows the estimated EPL gain covariance, corresponding to optimal hyperparameters *ρ*_*g*_ = 2.9, *ℓ*_*g*_ = 0.93, and *q* = 0.95.

**Figure 4.**
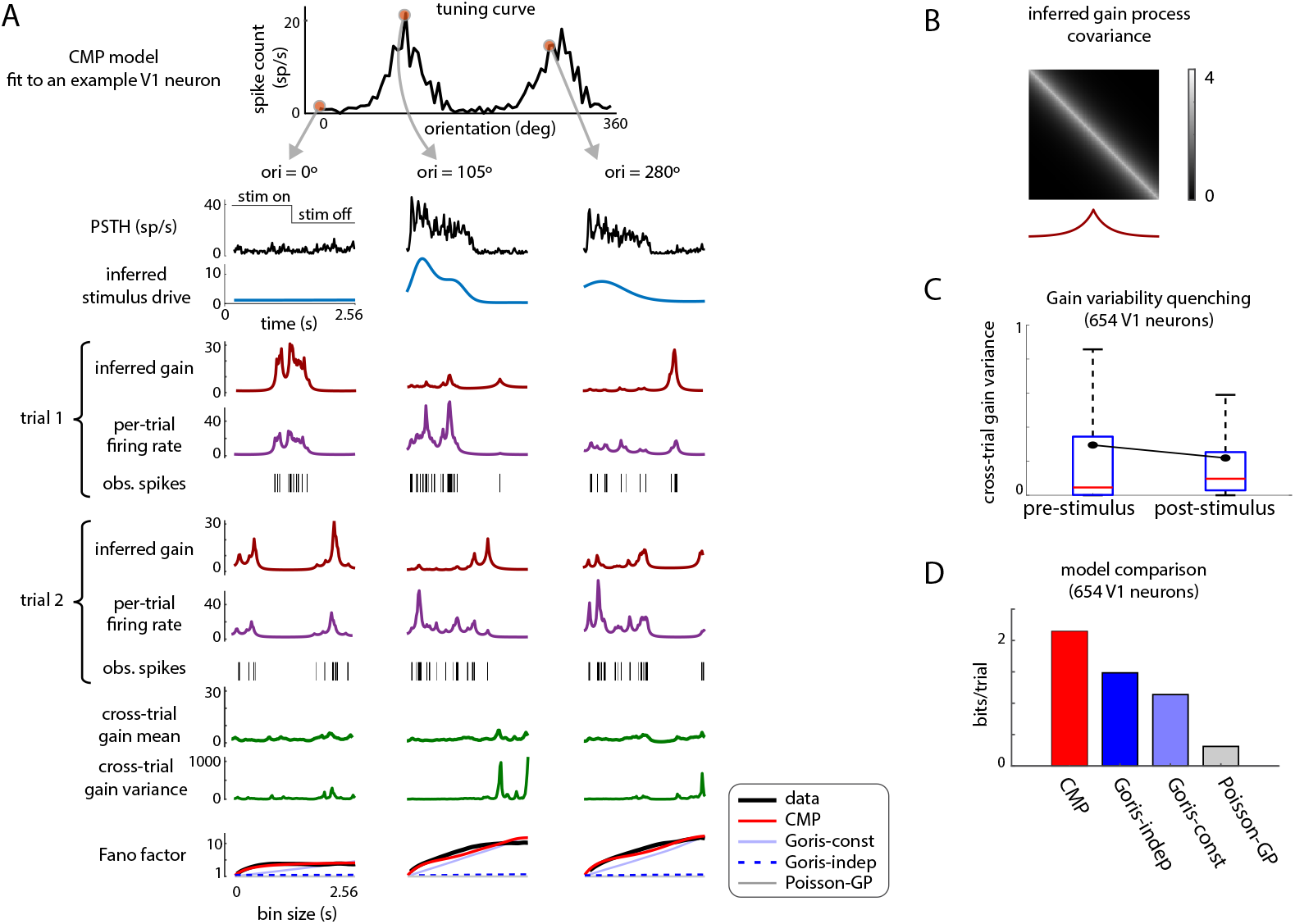
Performance comparisons of different models on neural population data recorded from macaque primary visual cortex under 72 drifting sinusoidal gratings (data from Graf et al. [36]). **(A)** Top to bottom: Tuning curve of a sample neuron; for three selected orientations (0^0^,105^0^,280^0^), the PSTH, inferred stimulus drive, inferred gain (trial 1), inferred firing rate (trial 1), observed spike raster (trial 1), inferred gain (trial 2), inferred firing rate (trial 2), observed spike raster (trial 2), cross-trial gain mean, cross-trial gain variance and Fano factor vs. bin size curves for real data (black), CMP model (red), Goris-independent gain model (dark blue dashed), Goris-constant gain model (light blue), and Poisson-GP model (gray). **(B)** EPL gain covariance (top) and autocorrelation (bottom) under the CMP model for the neuron in panel A. **(C)** Comparison of pre-stimulus and post-stimulus cross-trial gain variance across V1 neurons. Boxes show the distribution across neurons, and black dots indicate the mean. **(D)** Average test log-likelihood improvement of each model relative to the baseline Poisson model, computed on held-out data across all 654 V1 neurons and 72 stimulus orientations.

To further examine the temporal structure of inferred gain variability, we quantified cross-trial gain variance before and after stimulus onset. Following the approach of Churchland et al. [39], we compared gain variability in two matched 400-ms windows: a pre-stimulus window ending at stimulus onset and a stimulus-period window beginning 100 ms after stimulus onset. For each neuron and stimulus condition, we computed the cross-trial variance of the inferred gain at each time bin and averaged this quantity within each window. We then compared pre- and post-stimulus gain variability across neuron-stimulus pairs (Figure 4C). Across the 654 V1 neurons, post-stimulus gain variance was significantly lower than pre-stimulus gain variance (Figure 4C; one-sided paired Wilcoxon signed-rank test, *p* ≤ 10^−15^). This reduction is consistent with gain variability quenching following stimulus onset [39, 40].

The continuous-time formulation of the CMP model allows variability to be partitioned at arbitrary temporal resolutions, enabling analytical characterization of the relationship between overdispersion and bin size (see Methods and Materials for details). The bottom row of Figure 4A shows Fano factor curves derived from the CMP model (red), compared with curves computed from the data (black), the Goris model with independent gain (dark blue dashed), the Goris model with constant gain (light blue), and the Poisson-GP model (gray). The Poisson-GP model corresponds to the CMP model without a gain component, meaning that it uses an inhomogeneous Poisson observation model with a stimulus-specific, GP-smoothed time-varying firing rate. This differs from the baseline Poisson model, which estimates the time-dependent firing rate using a stimulus-conditioned PSTH with a Gamma prior for regularization in low-repeat conditions (see Methods and Materials for details of all model variants). The two variants of the Goris model are continuous-time extensions of the original modulated Poisson model [1]: the independent gain model assumes that the sampling interval of the gain process is much shorter than the trial duration, whereas the constant gain model assumes that the gain remains fixed throughout the entire trial. For both Goris variants shown here, the stimulus component was set to the GP-smoothed firing-rate estimate obtained from the Poisson-GP model, so that differences across these baselines reflect the assumed temporal structure of the gain process rather than differences in stimulus-drive estimation. These results show that the CMP model closely matches the overdispersion observed in real data across orientations and bin sizes, outperforming all other methods. While both Goris model variants produce overdispersion [1], the independent gain model yields Fano factor curves that are largely flat, and the constant gain model produces curves that increase linearly with bin size, neither of which captures the true empirical relationship.

We next compared model performance using the average log-likelihood improvement relative to the baseline Poisson model on held-out data from all 654 V1 neurons across all 72 orientations (Figure 4D). The CMP model achieved the highest predictive performance across all neurons. We also use these comparisons to separate the contribution of the GP-smoothed stimulus drive from the contribution of the stochastic gain process: the Poisson-GP model tests the effect of smoothing the stimulus-driven firing rate alone, whereas the CMP model additionally includes continuous-time gain fluctuations. In Figure S1, we further compared the CMP model to three variants with different gain kernels: CMP-RBF (where the EPL kernel in Equation 6 is replaced by the standard RBF kernel [33]), CMP-Matérn 0, and CMP-Matérn 1 (with Matérn kernels of order 0 and 1, respectively). These results demonstrate that the EPL kernel provides the best fit to the gain structure in the data.

To evaluate the utility of sharing gain hyperparameters across stimuli, we compared the CMP model with a variant in which the gain hyperparameters {*ρ*_*g*_, *ℓ*_*g*_, *q*} are fit independently for each of the *s* = 1, …, *S* stimulus conditions, denoted by 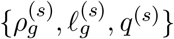. Figure S2A shows the inferred activity under both models, and Figure S2B presents the corresponding log-likelihoods. The performance of the two models was nearly identical, suggesting that the CMP model can robustly capture trial-to-trial variability across stimuli using a single shared set of gain hyperparameters.

Finally, we assessed the consistency of CMP inference with respect to temporal discretization by comparing inference results across three sufficiently fine grid resolutions: *dt* = 0.02 s, *dt* = 0.05 s, and *dt* = 0.1 s. As shown in Figure S3A, the estimated gain hyperparameters remain stable across these resolutions, indicating that the inference procedure is robust to moderate changes in grid size once the temporal discretization is fine enough to approximate the underlying continuous-time model. Figure S3B shows corresponding test log-likelihoods across models and grid resolutions, confirming that CMP consistently outperforms alternative models over this range.

### Continuous partitioning of variability along the visual hierarchy

Next, we evaluated model performance across different areas of the visual hierarchy by applying each method to neural responses recorded in the lateral geniculate nucleus (LGN), V1, V2, and MT (data from Goris et al. [1]). The recordings were obtained using drifting sinusoidal gratings of the preferred size and speed, varying either in *S* = 12 spatial frequency (ranging from 0 to 10 cycles/deg) or in *S* = 16 equally spaced drift direction. Each grating was presented for one second and repeated at least *N* = 5 times. We analyzed spiking activity from 48, 376, 167, and 127 neurons from LGN, V1, V2, and MT, respectively, using a grid resolution of *dt* = 0.02 s.

Figure 5A shows the average log-likelihood improvement on held-out data, relative to the baseline Poisson model, for each method across all four areas. The CMP model consistently outperformed all other models across the visual hierarchy. To examine performance at the single-neuron level, Figure 5B presents scatter plots comparing the log-likelihoods of the CMP model with those of the Poisson-GP model, the Goris independent gain model, and the Goris constant gain model. These results show that the CMP model systematically achieves better fits across individual neurons in all regions.

**Figure 5.**
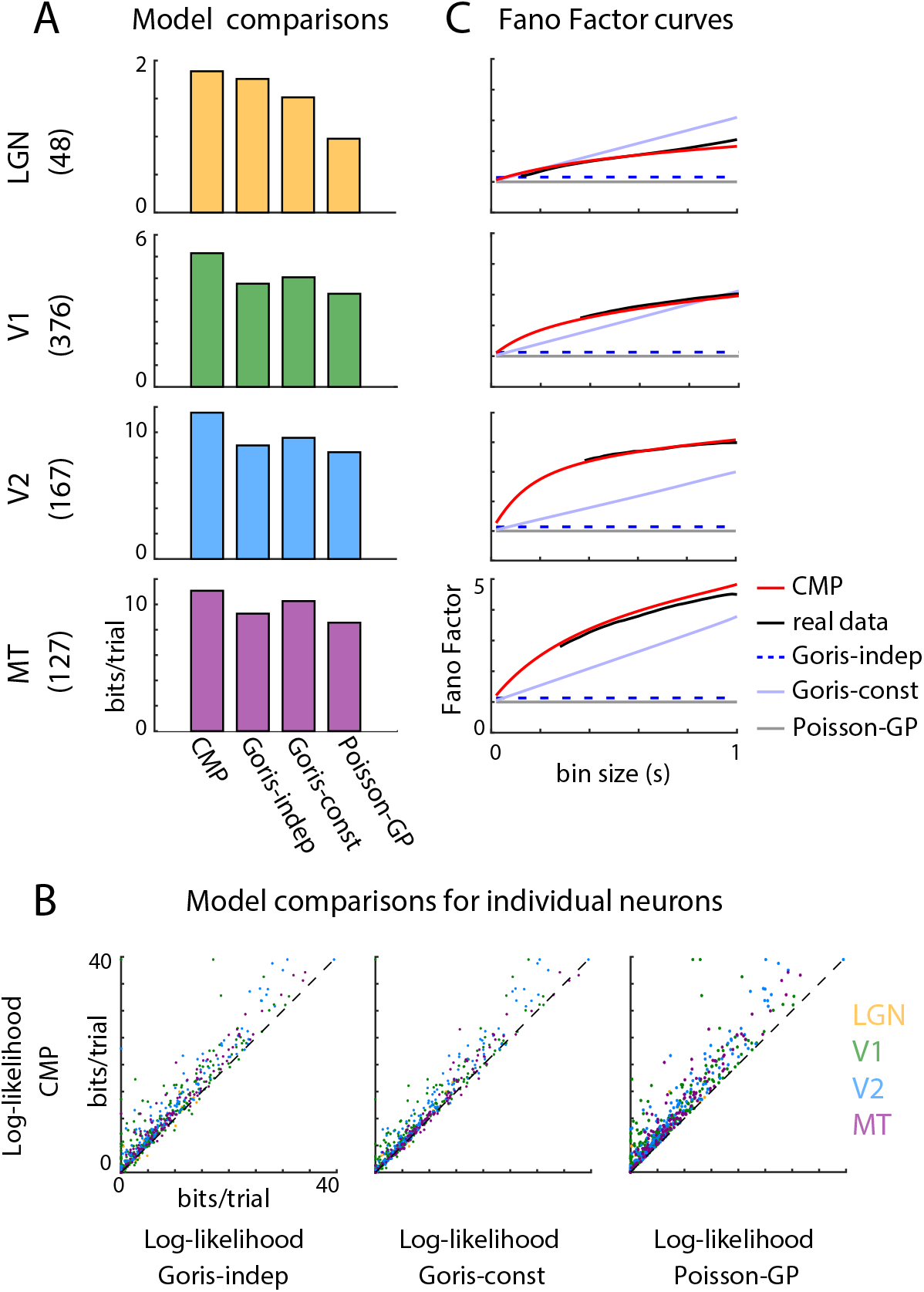
Performance comparisons of different models on neuronal population data recorded from different areas along the visual hierarchy—the lateral geniculate nucleus (LGN), V1, V2 and MT—under drifting sinusoidal gratings of preferred size and speed, varying in spatial frequency (12 frequencies from 0 to 10 cycles/deg) or in drift direction (16 directions) (data from Goris et al. [1]). **(A)** Log-likelihood improvement of each model relative to the baseline Poisson model, averaged across neurons in each population. Rows from top to bottom: LGN, V1, V2, MT. The number of neurons per area is indicated in parentheses. **(B)** Log-likelihood comparison of the CMP model versus the Goris independent gain model (left), Goris constant gain model (middle), and Poisson-GP model (right) for individual neurons. Colors indicate cortical areas: LGN (yellow), V1 (green), V2 (blue), MT (purple). **(C)** Fano factor vs. bin size curves averaged across each population. Black: real data; red: CMP; dark blue dashed: Goris-independent gain; light blue: Goris-constant gain; gray: Poisson-GP.

Next, we characterized trial-to-trial variability by analyzing the mean–variance relationship as a function of bin size. Figure 5C shows Fano factor curves, averaged over neurons in each population, for real data and the various models. Notably, the CMP model closely follows the overdispersion patterns observed in real data across visual areas and timescales, again outperforming alternative methods. In Figure S4, we extend this analysis by including three additional CMP variants: CMP-RBF, CMP-Matérn 0, and CMP-Matérn 1. These comparisons further confirm that the standard CMP model provides superior performance across all visual areas.

Finally, to examine the importance of including a GP prior on the stimulus drive, we compared the performance of the Goris-style models with and without this component (Figure S5). All Goris-style results reported above use versions with a GP prior on the stimulus drive, in which the stimulus-dependent firing rates were set to the smooth firing-rate estimates obtained from the Poisson-GP model, providing a stronger GP-smoothed stimulus-drive estimate for these baselines. In the versions without the GP prior, the stimulus-drive parameters were instead estimated directly under the corresponding Goris-style likelihood, making these models closer to the original modulated Poisson formulation [1] in their treatment of the stimulus drive. Figure S5 presents the log-likelihood comparisons from these analyses, which clearly show that incorporating a smoothness prior on the stimulus drive generally improves the model’s ability to fit held-out data. It is also noteworthy that the Poisson-GP model, despite lacking a stochastic gain component, outperforms both variants of the Goris-style model when the GP prior on the stimulus drive is removed (Figure S5A–C). These results indicate that the GP prior on the stimulus drive provides an important improvement in model fit, while the additional continuous-time gain component in CMP further improves the model’s ability to capture overdispersion and trial-to-trial variability (Figure S5D).

### Variations in gain and stimulus-drive along the visual hierarchy

Finally, we compared the distributions of the inferred CMP hyperparameters across the visual areas LGN, V1, V2, and MT, as shown in Figure 6. Panels A, B, and C show the distributions of the inferred gain power-law exponent *q*, gain length scale *ℓ*_*g*_, and gain variance *ρ*_*g*_, respectively, for each neuronal population. Figure 6D presents the median gain covariance and its corresponding autocorrelation function for each area.

**Figure 6.**
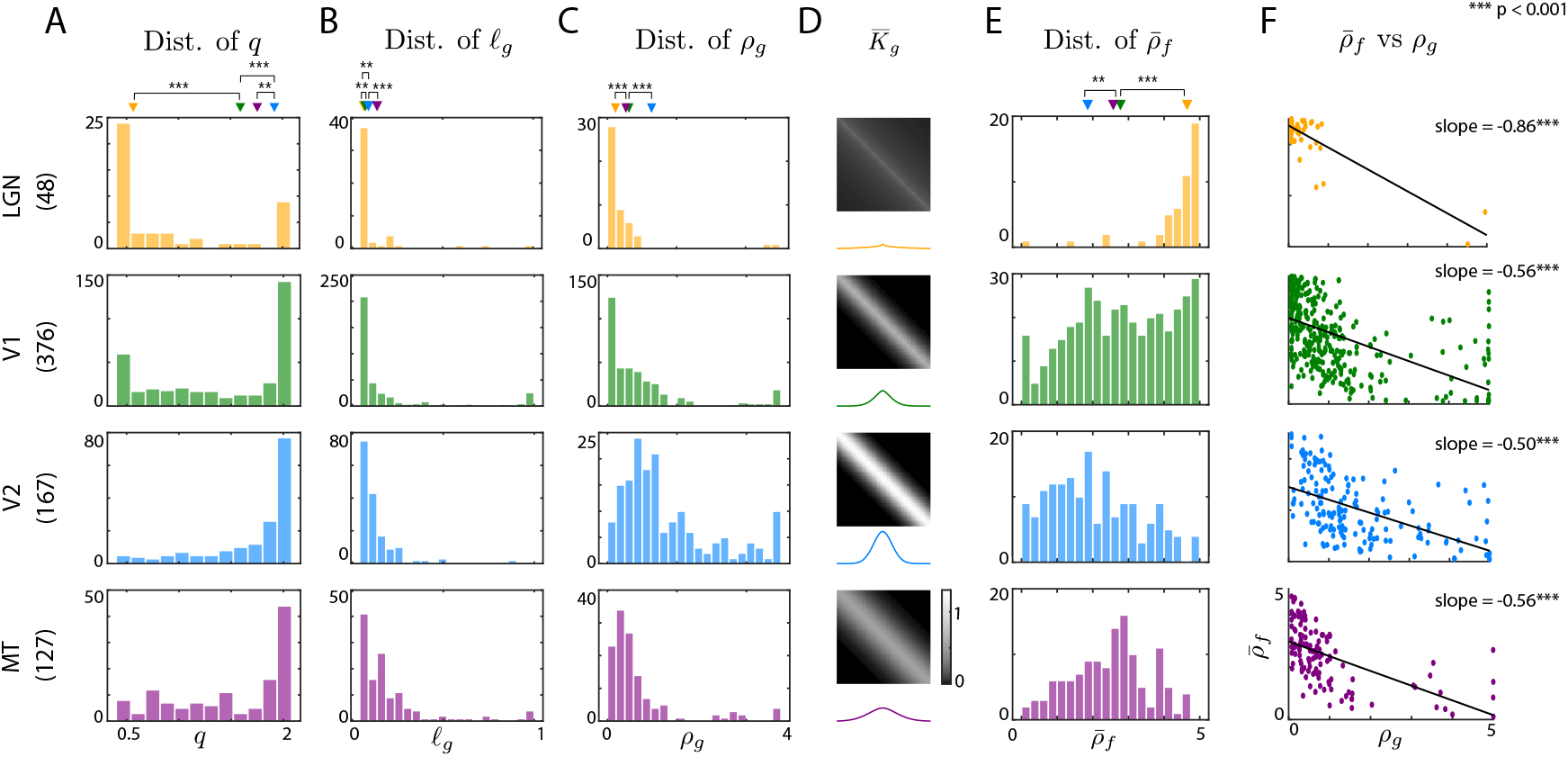
Comparison of inferred CMP hyperparameters across visual areas: the lateral geniculate nucleus (LGN), V1, V2, and MT (data from [1]). Rows top to bottom: LGN (yellow), V1 (green), V2 (blue), and MT (purple). The number of neurons per area is indicated in parentheses. **(A)** Distribution of gain power-law exponent *q* across neurons. Triangles indicate the median. **(B)** Distribution of gain length scale *ℓ*_*g*_ across neurons. **(C)** Distribution of gain variance *ρ*_*g*_ across neurons. **(D)** Median gain covariance 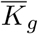 across the neuronal population (top) and gain auto-correlation (bottom). **(E)** Distribution of the average stimulus-drive variance 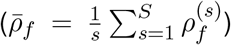 across neurons. **(F)** Scatter plots of 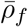 versus *ρ*_*g*_ across neurons, with linear regression fits (black) and slopes annotated. ^***^*p <* 0.001.

The gain power-law exponent (Figure 6A) increases along the visual hierarchy: from LGN to V1 (*p <* 0.001, sign test) and from V1 to V2 (*p <* 0.001), although there is a slight decrease from V2 to MT (*p <* 0.05). The gain length scale (Figure 6B) also shows a progressive increase: from LGN to V1 (*p <* 0.05), from V1 to V2 (*p <* 0.05), and from V2 to MT (*p <* 0.001). Additionally, the median gain autocorrelation (Figure 6D) exhibits slower temporal decay and a broader central lobe in higher visual areas. Together, these results suggest that rapid temporal fluctuations in trial-to-trial variability diminish as information propagates along the visual hierarchy.

In contrast, the gain variance *ρ*_*g*_ (Figure 6C) increases systematically along the hierarchy: from LGN to V1 (*p <* 0.001), and from V1 to V2 (*p <* 0.001). This trend indicates that the magnitude of gain fluctuations becomes larger at later stages of the visual pathway, consistent with prior observations [1].

Next, we examined the distribution of stimulus-drive variance, averaged across stimuli, defined as: 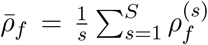, as shown in Figure 6E. A larger value of 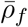 generally indicates stronger stimulus encoding by the neuron. In contrast to the gain variance trend, the stimulus-drive variance tends to decrease along the visual pathway: from LGN to V1 (*p <* 0.001), and from V1 to V2 (*p <* 0.05). This suggests a reduction in the strength of stimulus-driven encoding as information moves through the visual hierarchy.

To further investigate the relationship between stimulus-drive and gain variability, Figure 6F shows scatter plots of 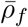 versus *ρ*_*g*_ for individual neurons in each area. Each panel includes a linear regression line with the slope indicated. Across all visual areas, there is a statistically significant negative correlation between the two quantities (*p <* 0.001, t-test), indicating that neurons with weaker stimulus-driven components tend to have stronger gain variability, and vice versa. This finding aligns with recent reports of an anti-correlation between stimulus encoding strength and trial-to-trial, stimulus-independent variability in sensory neurons [35, 41–43].

## 3 Discussion

Advances in high-density electrophysiological recording techniques [44, 45] now allow large-scale measurements of neuronal activity across multiple brain areas and experimental conditions. Analyses of sensory responses to repeated stimuli consistently reveal substantial trial-to-trial variability in neural spiking, arising from diverse sources such as synaptic transmission noise, fluctuations in excitation–inhibition balance, noisy peripheral inputs, recurrent dynamics, adaptation, and synaptic plasticity [6–11]. Capturing this variability is essential for understanding neural coding [12–14]. Traditional Poisson spiking models [18, 19] provide a foundational framework but predict spike count variance equal to the mean. This assumption is often violated in many brain regions where neurons exhibit overdispersion, meaning variability that exceeds the Poisson prediction [1]. Although numerous models have been proposed to account for this [23–29, 34, 35, 46, 47], few provide a detailed, time-resolved characterization of trial-to-trial variability.

Here, we introduced the Continuous Modulated Poisson (CMP) model, a principled probabilistic framework for modeling spiking variability in continuous time. The CMP model builds upon the modulated Poisson model [1] by describing the spike rate at each time point as the product of two independent processes: a stimulus-driven component and a stochastic gain process. Both components are modeled as exponentiated Gaussian processes (GPs), enabling flexible characterization of their smooth temporal dynamics. This model belongs to the class of log-Gaussian Cox processes [37], as it constitutes a doubly stochastic Poisson process. We allow the stimulus-driven components to have condition-specific hyperparameters, while the gain hyperparameters are shared across conditions, supporting efficient and interpretable joint inference.

To capture the long-range correlations observed in gain fluctuations, we introduced the Exponentiated Power Law (EPL) kernel, a new covariance function that generalizes the commonly used radial basis function (RBF) kernel. In our application, the EPL kernel consistently outperformed standard alternatives such as the RBF and Matérn kernels. In addition, we developed a scalable variational inference method that jointly estimates the latent stimulus-driven and stochastic gain components, along with their hyperparameters, directly from spiking data. We validated this approach using synthetic experiments. When applied to recordings from macaque V1 [36], the CMP model accurately partitioned response variability across time, trials, and stimulus conditions. It outperformed alternative approaches, including continuous-time extensions of the modulated Poisson model, in predicting held-out spiking activity and capturing the empirical Fano factor curves over time. Applying the CMP model to data from multiple stages of the visual hierarchy (LGN, V1, V2, and MT) [1], we again observed superior performance across areas and consistent predictions of overdispersion.

Analysis of the inferred CMP hyperparameters across visual areas revealed systematic trends. The temporal smoothness of gain fluctuations increased along the hierarchy, suggesting that trial-to-trial variability becomes more slowly varying in higher-order areas. In contrast, gain variance increased, consistent with an accumulation of shared noise or contextual modulation. Meanwhile, the variance of the stimulus-driven component decreased, consistent with increased selectivity and stabilization of tuning in downstream regions. A consistent negative correlation between the magnitudes of gain variance and stimulus-drive variance was observed across all areas, indicating that neurons with strong stimulus encoding tend to exhibit lower trial-to-trial variability, and vice versa. This observation aligns with previous findings of an anti-correlation between signal strength and variability in sensory neurons [35, 41–43]. These results have broader implications for understanding sensory coding, revealing how the brain balances fidelity and flexibility by modulating trial-to-trial variability across space and time [48]. They also align with broader theories in systems neuroscience, such as efficient coding [49] and probabilistic inference [50], where internal variability may be structured rather than random.

Our method complements black-box models by providing interpretable estimates of both stimulus driven and modulatory influences on spiking variability. It is particularly well-suited for experimental settings with repeated stimuli and temporally structured variability, such as studies of sensory tuning or motor control [1, 36, 51]. The CMP model preserves the interpretability of classical variability-partitioning approaches while extending them from spike-count models to a continuous-time spike-train framework. This enables time-resolved inference of stimulus-driven and stimulus-independent components, analytic characterization of Fano factor scaling with bin size, and direct estimation of the temporal covariance structure of gain fluctuations. While we found that the EPL kernel performed best in our experiments, kernel selection may depend on the task-specific temporal structure of neural variability.

More broadly, the EPL kernel introduced here is generally applicable and may support future Gaussian process–based modeling efforts beyond neuroscience. While the EPL kernel provides a flexible and tractable way to capture a wide range of temporal correlation structures in the stochastic gain process, its generative interpretation depends on the parameter *q*. For *q* = 1, a straightforward stochastic generator is a simple low-pass filtered Gaussian white noise process, equivalent to an Ornstein–Uhlenbeck (OU) process [52, 53]. For *q <* 1 there is no known closed-form stochastic differential equation that exactly produces the EPL covariance, but such long-memory dynamics can be closely approximated by superpositions of OU processes with a spectrum of time constants or by fractional stochastic processes such as fractional Brownian motion [54]. Conversely, when *q >* 1, correlations decay more rapidly than exponentially, corresponding to short-memory dynamics with diminished low-frequency power; these can often be approximated by higher-order linear filters or by driving OU processes with temporally correlated noise. Together, these interpretations highlight how the parameter *q* shapes the temporal structure of trial-to-trial variability.

An important direction for future work is to move beyond descriptive modeling and explore the biological interpretability of inferred hyperparameters. For instance, differences in *q* or *ℓ*_*g*_ across the visual hierarchy could reflect distinct circuit or cellular mechanisms, such as changes in synaptic integration timescales, recurrent dynamics, or bursting behavior. This perspective could help explain phenomena such as the disproportionately slow gain fluctuations we observe in the LGN (e.g., *q* ≈ 0.5), potentially linking them to specific biophysical or network-level processes. Establishing such connections would deepen our understanding of how structured variability arises and how it shapes sensory computation.

Mechanistic interpretations of structured variability may also benefit from parallels with other biological systems. For example, variability in bacterial flagellar motor switching and single-cell gene expression can be explained by a slow extrinsic process modulating a faster Poisson process, producing heavy-tailed dwell times or variance–mean scaling beyond Poisson expectations [55, 56]. This two-timescale structure closely parallels the CMP framework and suggests that future work incorporating explicit modulatory dynamics could offer deeper mechanistic insight into neural variability and its functional significance.

The present CMP framework captures Poisson and super-Poisson variability through a conditionally Poisson observation model with stochastic gain modulation. As a result, the model does not generate sub-Poisson spike count variability in its current form. Sub-Poisson variability has been observed in some neural systems and may arise from mechanisms such as refractoriness, spike-history dependence, regularized spiking, or other non-Poisson renewal structure [57–59]. Extending CMP to capture such effects would require modifying the observation model, for example by incorporating spike-history terms, renewal-process likelihoods, or alternative count distributions that allow variance to fall below the mean [13, 14, 29]. Such extensions could make it possible to partition both super-Poisson and sub-Poisson components of neural variability in continuous time and to examine how these components vary with behavioral state, attention, or arousal [60, 61].

Beyond these conceptual extensions, there are also several methodological directions for future research. First, extending CMP to high-dimensional population activity may allow joint modeling of shared variability across neurons. A natural direction would be to combine the CMP framework with ideas from Gaussian Process Factor Analysis [35] to introduce low-dimensional shared gain activity across neurons. Such an extension would allow CMP to capture correlated variability and population-wide modulatory dynamics in simultaneously recorded neural ensembles. Second, adapting the model to alternative link functions [26] could broaden its applicability to different statistical structures. Third, developing inference algorithms compatible with calcium imaging data [43] would extend its utility to optical recordings. Fourth, integrating behavioral covariates [62] or state variables such as attention [60] or arousal [61] could improve inference of nonstationary trial structure. Finally, while our focus here was on neural spike trains, the general formulation of the CMP model could be applied to other domains involving overdispersed count data [63–65].

To facilitate reproducibility and dissemination, we have released our implementation of the CMP model on GitHub [66]. In summary, the CMP framework provides a scalable, flexible, and interpretable approach for modeling neuronal spike variability, revealing new insights into the structure of trial-to-trial fluctuations and their organization across the visual hierarchy, while opening the door to deeper mechanistic interpretations of variability in neural systems.

## 4 Methods and Materials

### CMP model inference

We start by recalling the general CMP forward model:

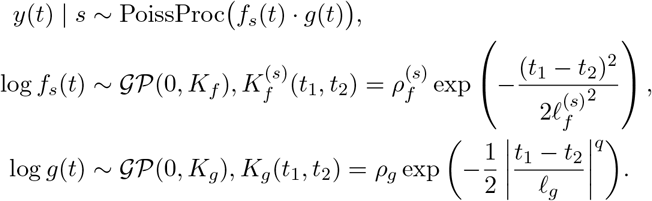

In our inference framework, we consider a setting with multiple repeats (*n* = 1, …, *N*) for a given stimulus condition, with data potentially collected across different stimulus conditions (*s* = 1, …, *S*). We perform inference on a uniform grid of time points (*t* = 1, …, *T*), with grid size set to *dt* seconds. Note that the model itself is defined in continuous time and can therefore be consistently applied to data measured at arbitrary temporal grid resolutions.

Let *y*_*t,s,n*_ denote the spike count for the *t*^th^ time bin, *s*^th^ stimulus condition, and *n*^th^ repeat. Further, let **f**_*s*_ = [*f*_1,*s*_ … *f*_*T,s*_]^⊤^ and **g**_*s,n*_ = [*g*_1,*s,n*_ … *g*_*T,s,n*_]^⊤^ denote the corresponding stimulus-driven and stochastic gain processes. Further, let 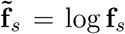 and 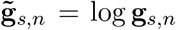 represent the log-transformed stimulus-driven and gain components, for notational brevity. Note that, following the CMP generative model, the latent variables 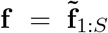 and 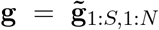, along with the hyperparameters 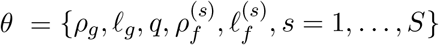, are all unknown quantities that need to be estimated given the spiking observations **y** = *y*_1:*T*,1:*S*,1:*N*_.

#### Frequency domain transformation of stimulus-driven components

The radial basis function (RBF) kernel diagonalizes in the Fourier domain, a property previously exploited to improve computational efficiency [35, 67]. Motivated by this, we adopt a frequency-domain representation of the stimulus-driven components (**f**) in our inference procedure. Accordingly, we relate the time-domain components to their corresponding Fourier-domain components:

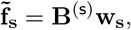

where 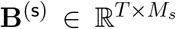 (noting that typically *M*_*s*_ ≪ *T* due to pruning of unnecessary Fourier coefficients that do not substantially contribute to explaining variability in 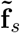 [35, 67]) represents the standard real orthonormal inverse discrete Fourier transform matrix. Then, with the diagonalization of the RBF kernel [35, 67], the Fourier-transformed stimulus-driven components 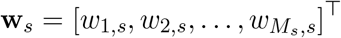 are distributed as independent Gaussian random variables with the following parameters:

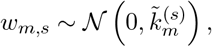

where 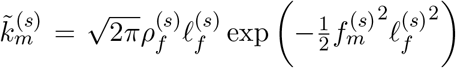, with 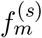 denoting the *m*^th^ Fourier frequency [35, 67]. Thus, we use the frequency-domain stimulus-driven components **w** = **w**_**1**:**S**_ in place of their time-domain counterparts **f**, reducing computational complexity by avoiding matrix inversions and enabling pruning (*M*_*s*_ ≪ *T*). We do not apply frequency-domain inference to the gain components **g**, as the heavy tails of the EPL kernel do not support effective pruning.

#### Variational Inference

To jointly infer the latent variables **w** and **g** and hyperparameters *θ*, we introduce an efficient inference procedure based on Variational Inference (VI) [38], which aims to find an approximate distribution *q* (**w, g**) for the intractable true posterior *p* (**w, g**|**y**, *θ*). In VI, the approximate posterior is first introduced into the data log-likelihood by rewriting it as [34]:

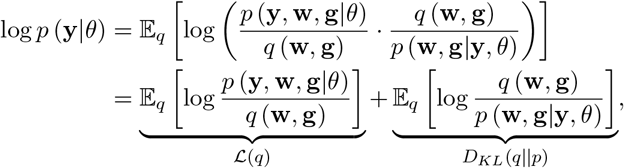

where *D*_*KL*_(*q*||*p*) is the Kullback-Leibler divergence between the variational approximation and the true posterior. Since *D*_*KL*_(*q*||*p*) is non-negative, ℒ(*q*) provides a lower bound on the model evidence, and is therefore called the Evidence Lower Bound (ELBO). Maximizing the ELBO is equivalent to minimizing *D*_*KL*_(*q*||*p*), thus finding the variational approximation closest to the true posterior.

#### Variational distributions and the ELBO

We assume that the variational distributions *q* (**w, g**) factorizes into independent Gaussian distributions over stimuli *s* = 1, …, *S* and trials *n* = 1, …, *N* :

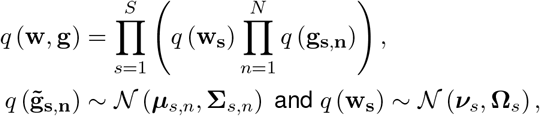

where **Σ**_*s,n*_ ∈ ℝ^*T* ×*T*^ and 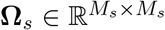 are full covariance matrices. Note that, by the property of linear transformations of Gaussian random variables [68], this in turn implies that the variational distributions over the time-domain stimulus-driven components are also Gaussian:

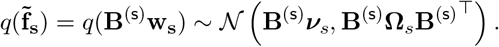

##### Algorithm 1

Joint inference of variational parameters and hyperparameters

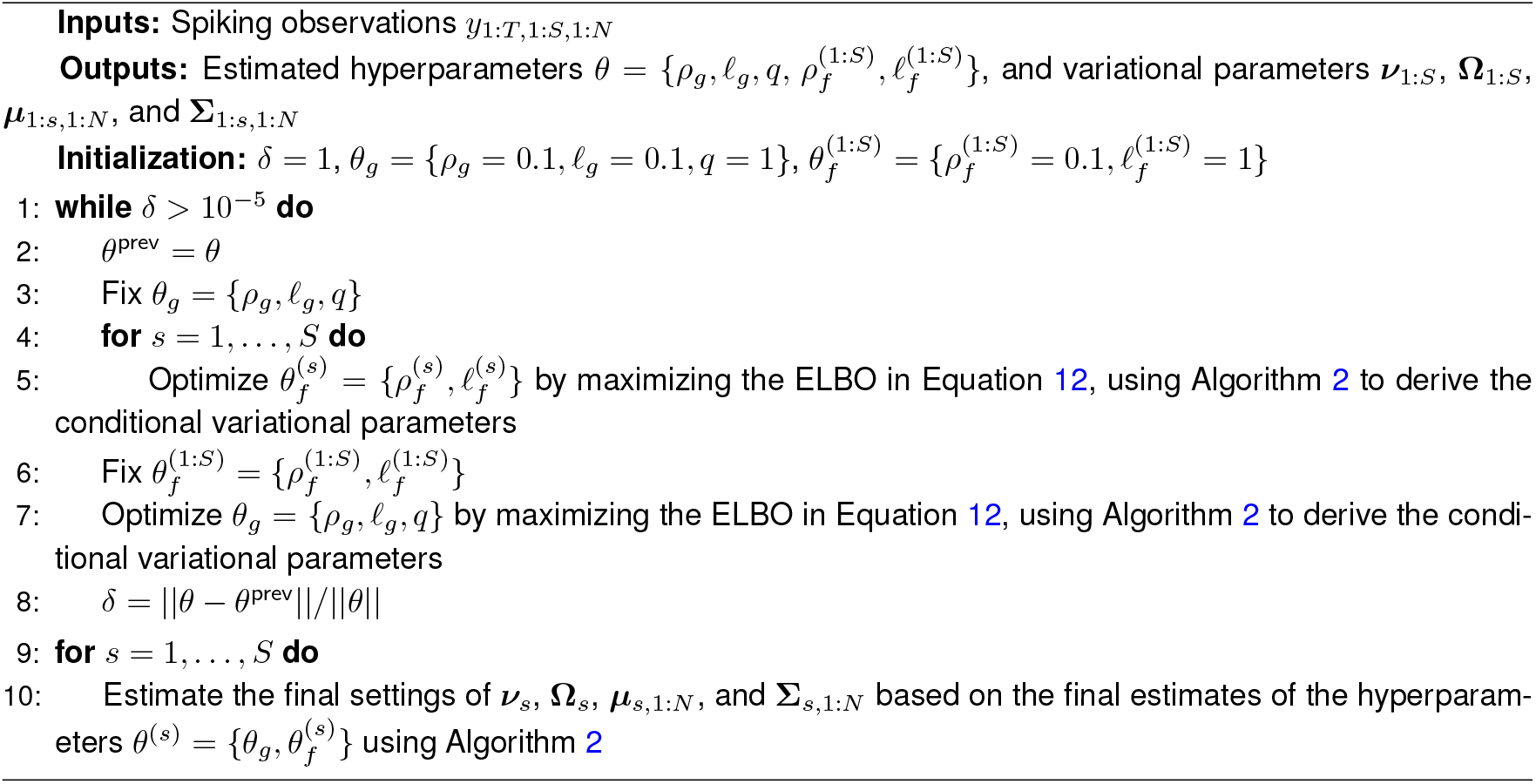

Accordingly, the ELBO can be derived as:

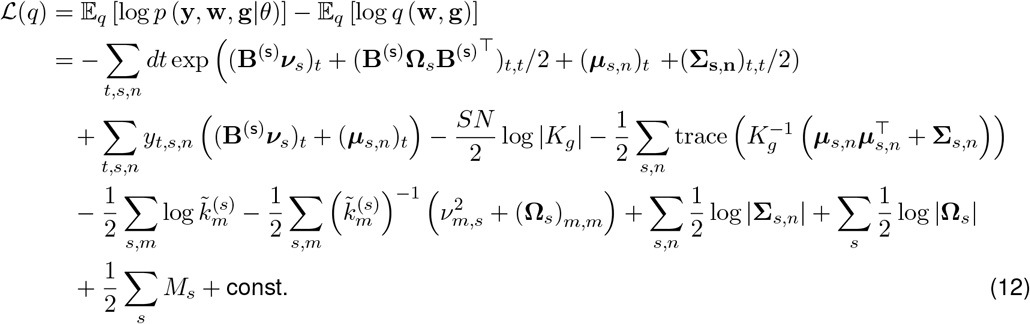

#### Joint inference of variational parameters and hyperparameters

In this section, we outline our proposed iterative scheme for the joint inference of variational parameters and hyperparameters, as illustrated in Algorithm 1. We alternate between optimizing the stimulus-drive hyperparameters 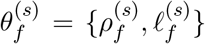 for *s* = 1, …, *S*, and the gain hyperparameters *θ*_*g*_ = {*ρ*_*g*_, *ℓ*_*g*_, *q*}, until convergence. Each optimization maximizes the ELBO in Equation 12, using Algorithm 2 (described in the next section) to infer the conditional variational parameters. We employed the built-in MATLAB function *fmincon*, a gradient-based solver for constrained nonlinear multivariable problems, to perform each maximization. To ensure stability, convergence, and valid covariance matrices, we constrained the hyperparameters to the following ranges:, 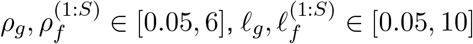 and *q* ∈ [0.4, 2].

##### Algorithm 2

Estimation of variational parameters given hyperparameters

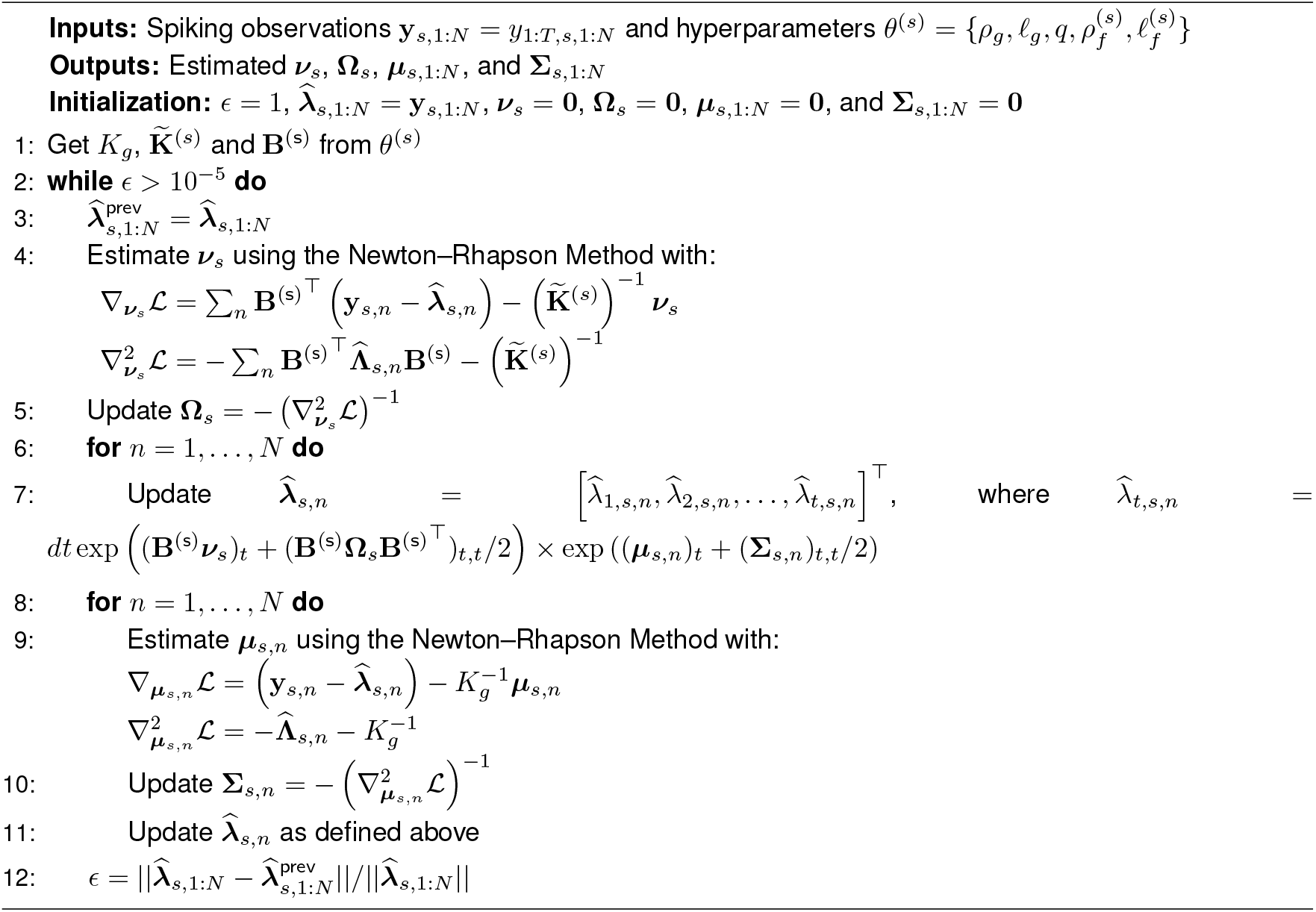

#### Inferring variational parameters given hyperparameters

Here, we outline the procedure for inferring the variational parameters by optimizing the ELBO for a given set of hyperparameters *θ* (i.e., assuming *K*_*g*_ and 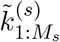 are known), as summarized in Algorithm 2. Note that the ELBO in Equation 12 is independent across stimulus conditions *s* = 1, …, *S*, when conditioned on *θ*. Therefore, we focus on estimating the variational parameters corresponding to the *s*^th^ stimulus: ***ν***_*s*_, **Ω**_*s*_, ***µ***_*s*,1:*N*_, and **Σ**_*s*,1:*N*_. We maximize the ELBO to find the optimal settings of these variational parameters, alternating between updating the stimulus-drive variational parameters (***ν***_*s*_, **Ω**_*s*_) and the gain variational parameters (***µ***_*s*,1:*N*_, **Σ**_*s*,1:*N*_) until convergence. Each maximization step uses a Newton–Raphson procedure.

First, consider the estimation of the stimulus-drive variational parameters ***ν***_*s*_ and **Ω**_*s*_, given the estimated gain variational parameters from the previous iteration. The gradient and Hessian of Equation 12 with respect to ***ν***_*s*_ can be derived as:

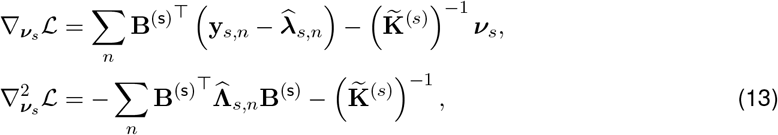

where 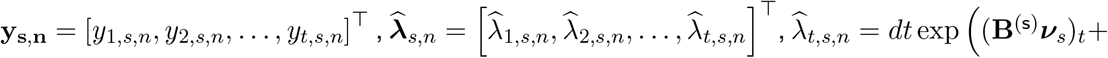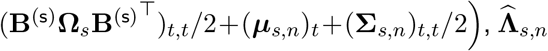 is a diagonal matrix with the *t*^th^ diagonal entry given by 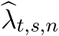, and 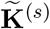 is a diagonal matrix with the *m*^th^ diagonal entry equal to 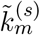. Further, taking derivatives in Equation 12 with respect to **Ω**_*s*_ yields:

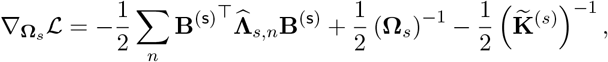

and setting this derivative to zero results in:

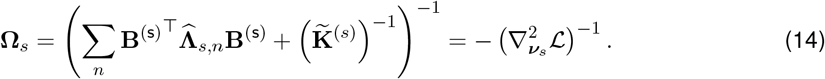

Accordingly, we update the stimulus-drive variational mean ***ν***_*s*_ using Newton–Raphson optimization with the gradient and Hessian in Equation 13, and subsequently update the stimulus-drive variational covariance **Ω**_*s*_ using the Hessian as in Equation 14.

Next, we update the gain variational parameters independently for *n* = 1, …, *N* using the updated stimulus-drive variational parameters, following an equivalent Newton–Raphson procedure, with the gradient and Hessian relative to ***µ***_*s,n*_ given by:

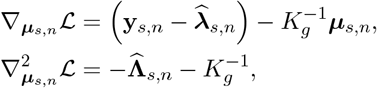

and the update rule of **Σ**_*s,n*_ given by:

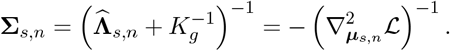

Following the above derivations, we iterate between updating the stimulus-drive variational parameters and gain variational parameters until convergence. The combined iterative procedure for variational parameter inference is summarized in Algorithm 2.

### Log-likelihood derivation of the CMP model

While we optimize the ELBO for parameter inference, accurately and fairly comparing models requires computing the log-likelihood on held-out data. To estimate this for the CMP model, we used the importance sampling procedure described below. Note that,

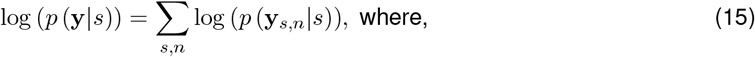

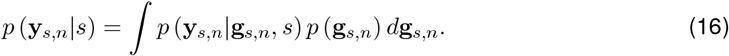

Observing that the integral in Equation 16 is intractable, we used importance sampling [69] to approximate it. We employed the variational distribution 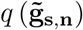 as the proposal distribution,

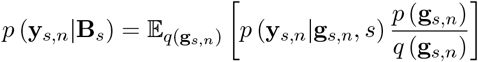

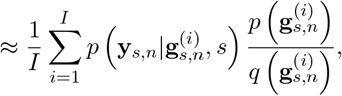

where 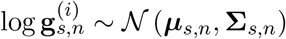. Defining:

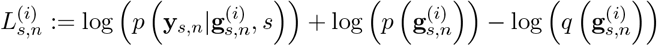

and applying the log-sum-exp trick for numerical stability, we obtained:

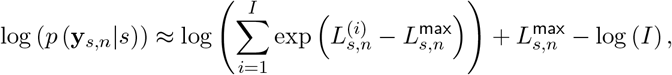

where 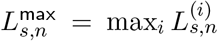. We set *I* = 1000 and used the above formulation to estimate the log-likelihood of the spiking observations in the *n*^th^ test trial of the *s*^th^ stimulus, and finally derived the joint log-likelihood of all test data according to Equation 15.

### Partitioning variability with CMP

Here we provide the general formula for partitioning variability into a Poisson component, equal to the mean, and a modulatory component, based on the CMP model in the setting where the stimulus drive *f*_*s*_(*t*) is a time-varying signal. The continuous-time formulation of the CMP model allows this partitioning to be computed for arbitrary time windows, enabling an analytic characterization of how overdispersion grows with the bin size used to count spikes. For a time bin of size Δ, the mean is given by:

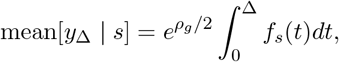

and the variance is:

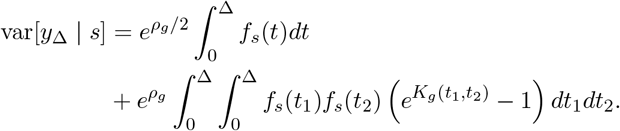

From these expressions, we can compute the Fano factor as a function of bin size, which is simply the ratio of variance to mean:

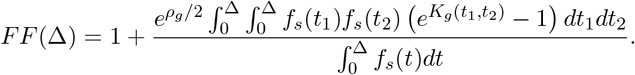

The details of the derivation are provided in the Appendix 2. To summarize statistics at a given bin size Δ over the entire observation window *T*, we use the following approach: we apply a sliding window of duration Δ, shift it systematically through the entire duration *T*, and average the computed statistics across all such windows.

### Alternative models considered for performance comparison

We compared the performance of the CMP model with the alternative models described below.

#### Baseline Poisson

All log-likelihood figures reported in this paper are expressed as log-likelihood gains of each method relative to the baseline Poisson model. The baseline Poisson model assumed spike counts across trials are generated by a Poisson process:

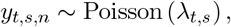

and, to compensate for sparsity, we further assumed a Gamma prior on the firing rate *λ*_*t,s*_ ~ Gamma (*α, β*), where *α* and *β* are hyperparameters. Since the Gamma distribution is conjugate to the Poisson distribution [70], the posterior distribution is also Gamma-distributed:

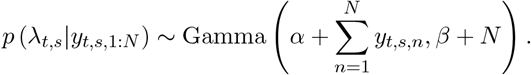

Thus, we found the MAP estimate of the firing rate by the mode of the posterior Gamma distribution, given by:

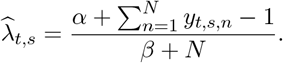

To fix the hyperparameters, we performed moment matching with the mean 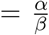 and variance 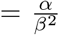. Note further that the mean and variance conditioned on *λ*_*t,s*_ are identical for this model, resulting in a Fano factor of 1.

#### Poisson-GP

This model includes only the stimulus-driven components, assuming that trial-to-trial variability is fully described by the stochastic nature of the Poisson process:

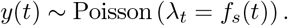

The stimulus drive itself is modeled by a Gaussian process, similar to the CMP model:

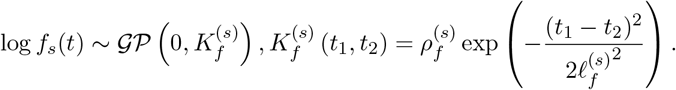

We inferred the hyperparameters and latent stimulus-driven components of this model using a simplified version of our proposed variational inference procedure. As in the baseline Poisson model, the mean and variance conditioned on the stimulus are identical, yielding a Fano factor of 1.

#### Goris-indep

This model extends the modulated Poisson framework of Goris et al. [1] to continuous time. The model in Goris et al. [1] considered spike counts within a single time bin. For spiking observations from the *s*^th^ stimulus presentation with *t* = 1, …, *T* time bins, we assumed that the stimulus-independent gain is independent across time bins with shared variance 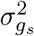, i.e.,

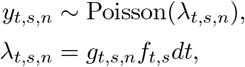

where *g*_*t,s,n*_ ~ Gamma(*r*_*s*_, *s*_*s*_), with shape parameter 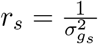 and scale parameter 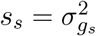. Marginalizing over the gain variables yields a product of independent Negative Binomial distributions:

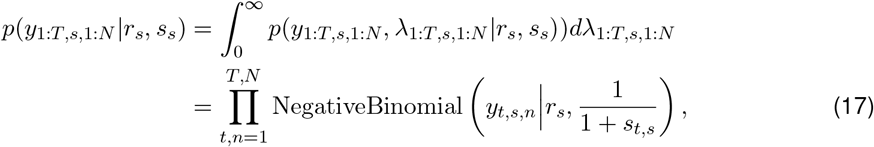

where 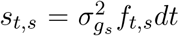. For the Goris-indep baseline used in the main model comparisons, we set the stimulus component to the smooth firing-rate estimate obtained from the Poisson-GP model. Specifically, we set 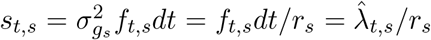, where 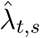 denotes the Poisson-GP estimate of the stimulus-dependent firing rate. For *s* = 1, …, *S*, we searched for the values of *r*_*s*_ that minimize the negative log-likelihood of the spiking observations *y*_1:*T,s*,1:*N*_. Details of Equation 17, the Fano factor, and the version without a smoothness prior on the stimulus drive are provided in the Appendix 3.

#### Goris-const

This is another extension of the Goris model [1] to continuous time. Here, we assumed the gain is constant across time bins within a trial:

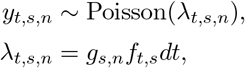

where *g*_*s,n*_ ~ Gamma(*r*_*s*_, *s*_*s*_) with shape parameter 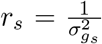 and scale parameter 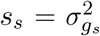. Further, we assumed that the total spike count across all time bins in a trial, *Y*_*s,n*_ = ∑_*t*_ *y*_*t,s,n*_,, also shares the same gain variance 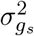, i.e, 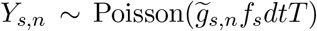 with 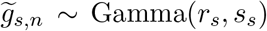. For the Gorisconst baseline used in the main model comparisons, we estimated 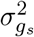 by fitting *Y*_*s,n*_ across training data *n* = 1, …, *N*, as in Goris et al. [1], and set the stimulus encoding component 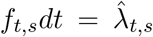, where 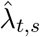 denotes the smooth firing-rate estimate obtained from the Poisson-GP model. Details of the mean-variance relationship, log-likelihood derivation, and the version without a smoothness prior on the stimulus drive are provided in the Appendix 3.

#### CMP-RBF

This model follows the same generative structure as the CMP model, except that the gain covariance is parameterized by the standard RBF kernel:

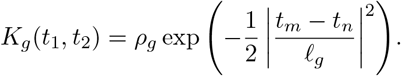

#### CMP-Matérn

This variant also follows the CMP framework, with the gain covariance parameterized by a Matérn kernel of order 0 in the CMP-Matérn-0 model:

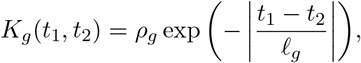

and a Matérn kernel of order 1 in the CMP-Matérn-1 model:

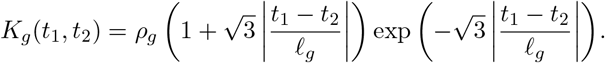

We used the same variational inference procedure as the CMP model to infer the hyperparameters and latent variables of the CMP-RBF and CMP-Matérn models. The log-likelihood and mean-variance derivations for these models are analogous to those of the CMP model.

### Datasets studied

In this paper, we applied the CMP model to two datasets recorded from anesthetized, paralyzed, adult macaque monkeys.

#### Multi-electrode array recordings from the primary visual cortex

Complete details about this dataset are available in [36]. In brief, extracellular recordings were obtained using a 10-by-10 fixed silicon microelectrode array (length 1mm, spaced 400 *µ*m) implanted in the superficial layers of the primary visual cortex (area V1) of three adult male macaque monkeys. The animals were stimulated with drifting sinusoidal gratings at full contrast, with the orientation of the grating varied around the clock in steps of 5 degrees, yielding a total of 72 orientations. Each stimulus was presented for 1,280ms and followed by a blank screen of mean luminance for another 1,280ms, repeated for 50 trials per stimulus condition. We analyzed 654 isolated units from five datasets in this paper (147, 113, 113, 133, and 148 units in each dataset). For each unit and each stimulus, we randomly selected *N* = 45 trials for training (i.e., for hyperparameter and stimulus-drive inference) and used the remaining *N* = 5 trials to derive the test log-likelihood.

#### Single-electrode recordings along the visual hierarchy

Full details regarding these datasets are available in [1]. Briefly, extracellular recordings were made from 18 adult macaque monkeys of either sex, in different areas along the visual hierarchy: the lateral geniculate nucleus (LGN), V1, V2, and MT. The animals were presented with drifting sinusoidal gratings of the preferred size and speed, varying either in spatial frequency (12 spatial frequencies, ranging from 0 to 10 cycles/deg) or in drift direction (16 equally spaced directions). Each grating was presented for 1,000 ms and repeated at least five times. We analyzed the activity of 48, 376, 167, and 127 cells from each population, respectively. For each unit and each stimulus, we randomly selected *N* = 1 trial to derive the test log-likelihood and used the remaining trials for training the model.

## 5 Acknowledgments

We are grateful to Arnulf Graf, Robbe Goris, Eero Simoncelli, and J. Anthony Movshon for providing the datasets used in this study. AR was supported by NIH T32 institutional training grant (T32MH065214). ASC was supported by NSF CAREER award 2340338. JWP was supported by grants from the Simons Collaboration on the Global Brain (SCGB AWD543027) the NIH BRAIN initiative (R01DA056404, and R01EB02694), the National Eye Institute (NEI) of the National Institutes of Health (NIH) (R01EY033064), and a U19 NIH-NINDS BRAIN Initiative Award (5U19NS123716).

## Appendix 1 Proof of the validity of the EPL covariance function

In this section, we provide the formal proof that the proposed Exponetiated Power Law (EPL) kernel is a valid covariance function.

### Theorem 1

(Validity of the EPL Kernel). *The Exponetiated Power Law (EPL) kernel given by:*

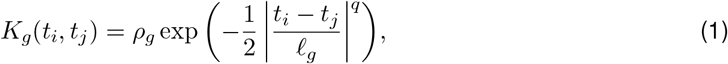

*where ρ*_*g*_ *>* 0 *is the marginal variance, ℓ*_*g*_ *>* 0 *is the length scale, and* 0 *< q* ≤ 2 *is the power law exponent, is a valid covariance function*.

*Proof*. We begin by recalling the definition of a valid covariance function [1].

### Definition 1

(Covariance function). *A (real-valued) function*

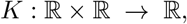

*is called a* covariance kernel *(or* positive semidefinite kernel*) if, for every finite set of points t*_1_, …, *t*_*n*_ ∈ ℝ *and all real weights a*_1_, …, *a*_*n*_ ∈ ℝ, *the following Gram sum is non-negative:*

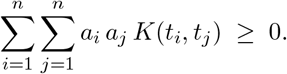

We now invoke the following result from the Schoenberg’s theorem [2].

### Lemma 1

(Schoenberg (1938)). *The function* exp (−|*x*|^*q*^) *is positive definite on* ℝ *for all* 0 *< q* ≤ 2. *Proof of Lemma 1*. The proof is provided in [2].

From Lemma 1 and Definition 1, it follows that for any finite set 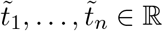 and weights *a*_1_, …, *a*_*n*_ ∈ ℝ,

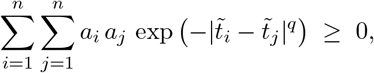

if 0 *< q* ≤ 2.

Next, consider the change of variables 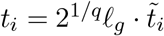, which implies:

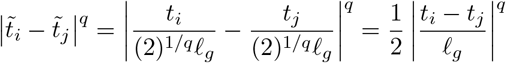

Substituting this into the above expression, we obtain:

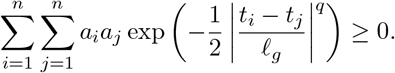

Finally, since *ρ*_*g*_ *>* 0, multiplying by *ρ*_*g*_ preserves non-negativity:

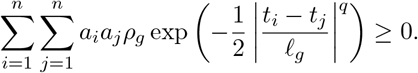

Hence, for any finite set *t*_1_, …, *t*_*n*_ ∈ ℝ and any real weights *a*_1_, …, *a*_*n*_ ∈ ℝ, we have:

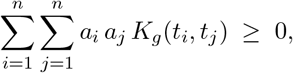

which establishes that the EPL kernel in Equation 1 is positive semidefinite, and thus a valid covariance function.

## Appendix 2 Partitioning variability with CMP

In this section, we provide the detailed derivations of the mean–variance relationship under the CMP model. Under the CMP model, spike train variability can be partitioned into a Poisson component, which is equal to the mean, and a modulatory component, which controls the degree of overdispersion. The continuous-time formulation of the CMP model allows us to compute this partitioning for arbitrary time intervals (Δ), and thus to analytically characterize how overdispersion grows with time bin size.

Recall the CMP forward model:

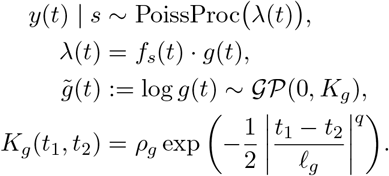

Under this model, the mean spike count in a time bin of size Δ, conditioned on the stimulus *s*, can be derived as:

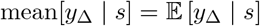

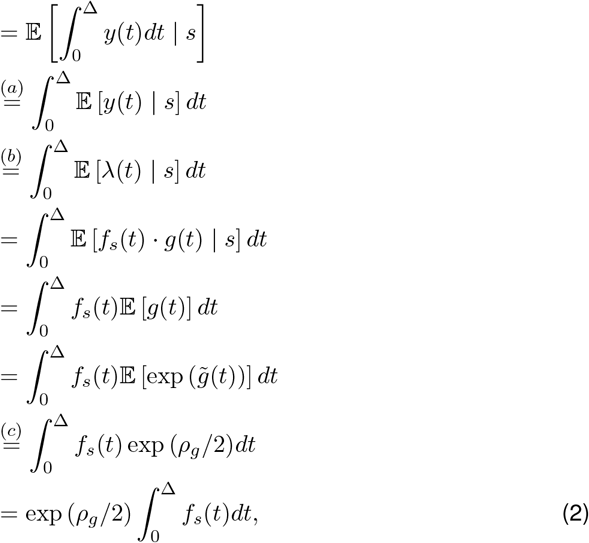

where (*a*) follows from Fubini’s Theorem [3], (*b*) uses the fact that the mean of a Poisson process equals its rate function [3], and (*c*) applies the moment-generating function of a Gaussian (i.e., the mean of a log-normal distribution) [4].

Using similar steps, the second moment can be derived as:

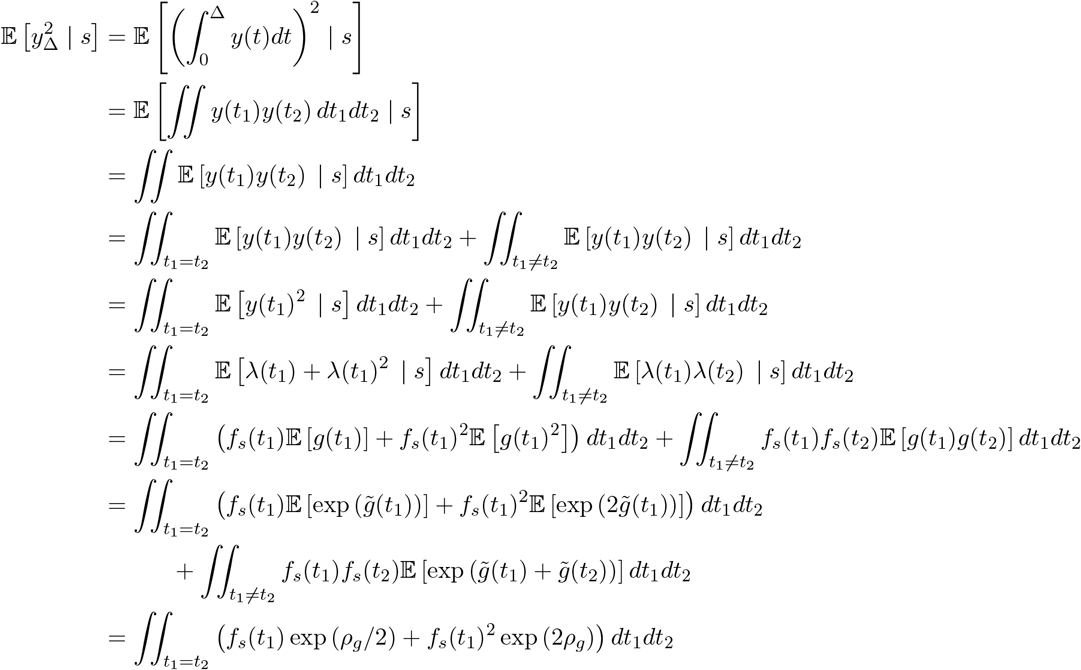

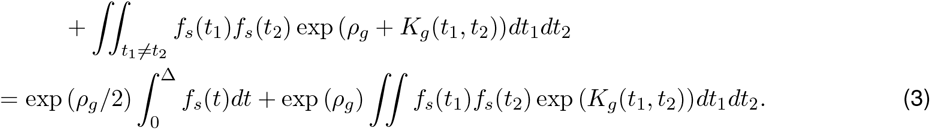

Thus, using Equation 2 and Equation 3 the variance can be formulated as:

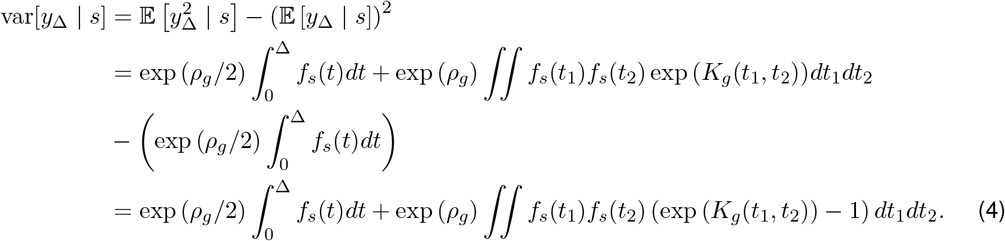

From these expressions, we can compute Fano factor as a function of bin size, which is simply the ratio of variance (Equation 4) to mean (Equation 2):

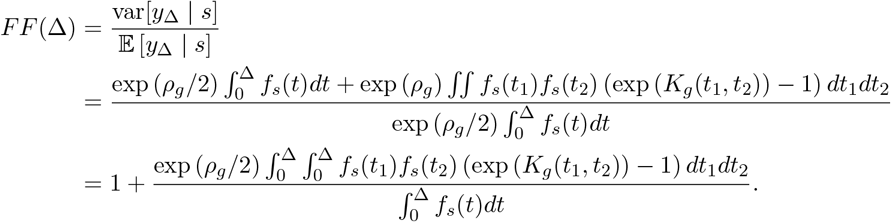

In the special case where the stimulus component is non-fluctuating, i.e., 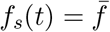, these expressions simplify to:

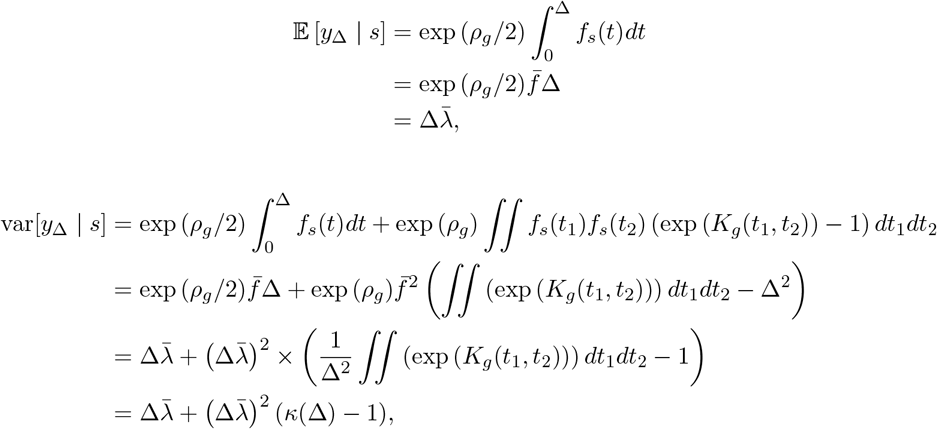

and:

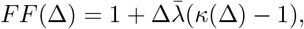

where we define 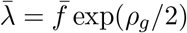 and 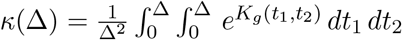.

## Appendix 3 Details about the Goris model variants

In this appendix, we provide technical details for two extensions of the modulated Poisson framework introduced by Goris et al. [5], adapted for spike-train data with multiple time bins per trial. Specifically, we derive the likelihoods, means, and variances for the Goris-indep model, which assumes time-bin–independent modulatory gains, and the Goris-const model, which assumes a constant gain across time bins within a trial. These derivations clarify the statistical properties of each model and describe how we compute log-likelihoods for model comparison. All Goris-style baselines reported in the main model comparisons use a smooth stimulus-drive estimate obtained from the Poisson-GP model, so that comparisons among these baselines focus on the assumed temporal structure of the gain process. We also evaluate versions without this smoothness prior, in which the stimulus-drive parameters are estimated directly under the corresponding Goris-style likelihood. Details for both versions are provided below.

### Appendix 3.1 Goris-indep model

Here we provide further details on the Goris-independent gain model, including the derivation of the likelihood, as well as the mean and variance under this model. The Goris-indep model is an extension of the modulated Poisson model in Goris et al. [5], adapted for spike train data with multiple time bins per trial. Given spiking observations of the *s*^th^ stimulus presentation across *t* = 1, …, *T* time bins, we assume that the modulatory gain process is independent across time bins, with shared variance 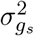, i.e.,

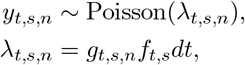

where *g*_*t,s,n*_ ~ Gamma(*r*_*s*_, *s*_*s*_), with shape parameter 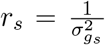 and scale parameter 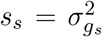. This implies *λ*_*t,s,n*_ ~ Gamma (*r*_*s*_, *s*_*t,s*_), where 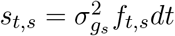.

#### Appendix 3.1.1 Likelihood derivation

Following the results in [5], marginalizing over the gain variable yields independent Negative Binomial distributions:

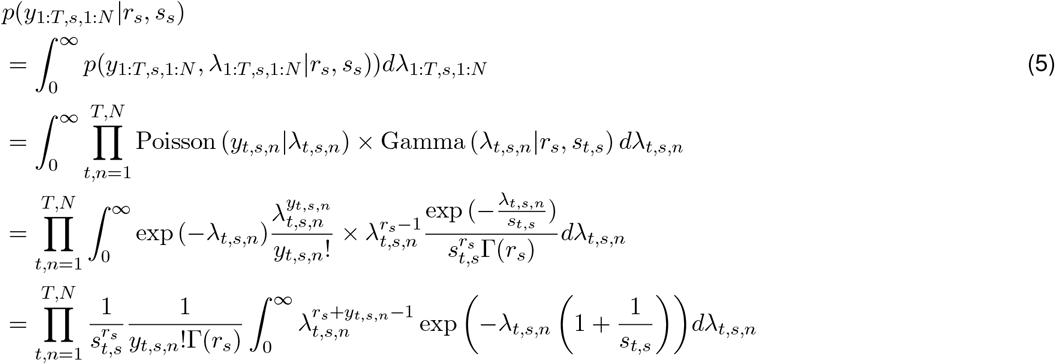

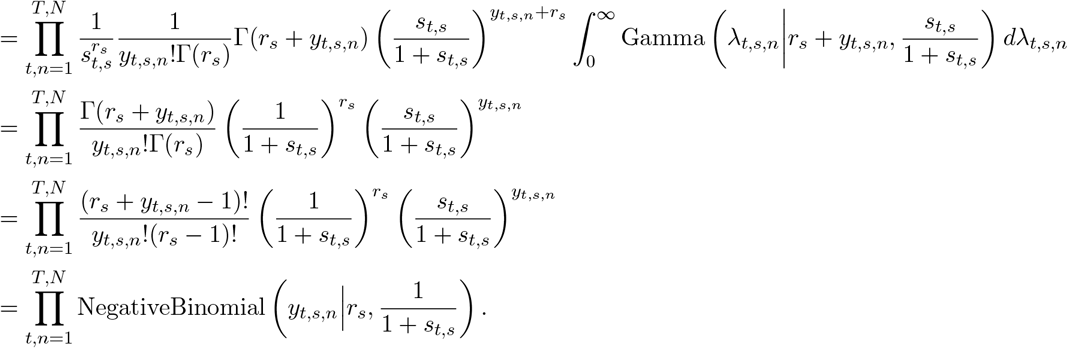

For the Goris-indep model used in the main model comparisons, we set the stimulus component to the smooth firing-rate estimate obtained from the Poisson-GP model. Specifically, we set:

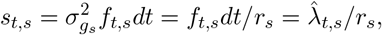

where 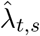 denotes the Poisson-GP estimate of the stimulus-dependent firing rate. We then search for the value of *r*_*s*_ that minimizes the negative log-likelihood of the spiking observations *y*_1:*T,s*,1:*N*_ for each stimulus condition. For the comparison without a smoothness prior on the stimulus drive, we instead estimate both *r*_*s*_ and *s*_1:*T,s*_ directly by minimizing the negative log-likelihood under the above Negative Binomial mixture model, as in Goris et al. [5].

#### Appendix 3.1.2 Mean-variance derivations

Based on the above derivation, we see that:

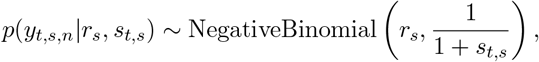

and *y*_1:*t,s,n*_ are independent across *n* = 1, …, *N* when conditioned on *r*_*s*_ and *s*_*t,s*_. Thus, from the mean and variance of the Negative Binomial distribution, we have:

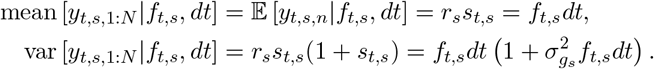

Next, we extend this to derive the mean and variance of spike counts over a larger time window Δ = *R dt*, by summing across the relevant bins *t*_1_, …, *t*_*R*_:

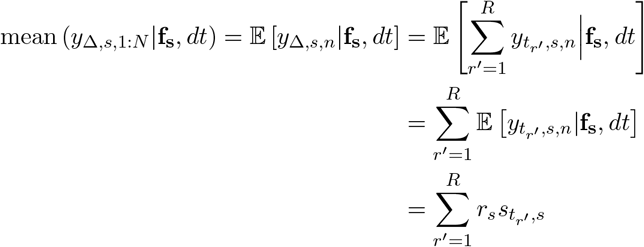

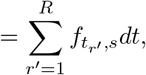

and:

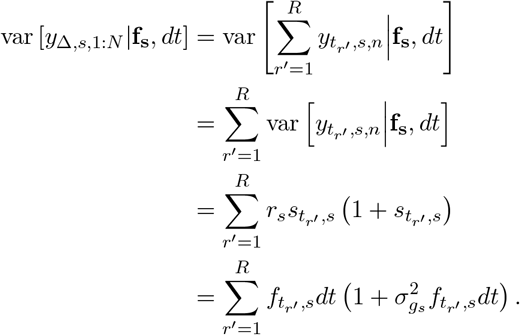

### Appendix 3.2 Goris-const model

In this section, we provide further details on the Goris-constant gain model, including the derivation of the mean and variance, as well as the log-likelihood under its modeling assumptions. The Goris-const model is the second variant we consider for extending the Goris model in [5] to multiple time bins. Here, we assume that the modulatory gain process is constant across time bins:

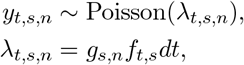

where *g*_*s,n*_ ~ Gamma(*r*_*s*_, *s*_*s*_), with shape parameter 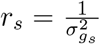 and scale parameter 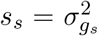. Furthermore, we assume that the total spike count across all time bins in a trial, *Y*_*s,n*_ = ∑_*t*_ *y*_*t,s,n*_, shares the same gain variance 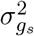, i.e.,

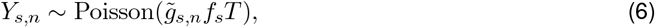

with 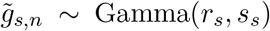. For the Goris-const model used in the main model comparisons, we estimate 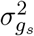 by fitting the total spike counts *Y*_*s,n*_ across training trials, as in Goris et al. [5], and set the stimulus encoding component 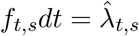, where 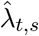 denotes the smooth firing-rate estimate obtained from the Poisson-GP model. For the comparison without a smoothness prior on the stimulus drive, we instead estimate *f*_*t,s*_*dt* using the same method as in the baseline Poisson model.

#### Appendix 3.2.1 Mean-variance derivations

Note that from the moments of the Gamma distribution:

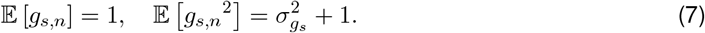

Accordingly, we derive the first moment, second moment, and variance at bin size *dt* as:

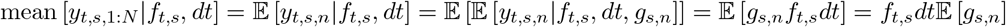

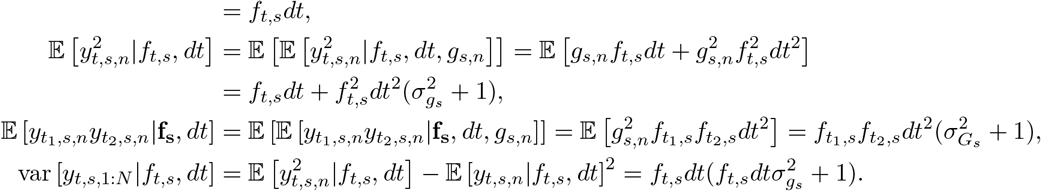

We then extend these results to a larger bin size Δ = *R dt*, by integrating over the differential statistics of the corresponding time bins *t*_1_, · · ·, *t*_*R*_ at bin-size *dt*:

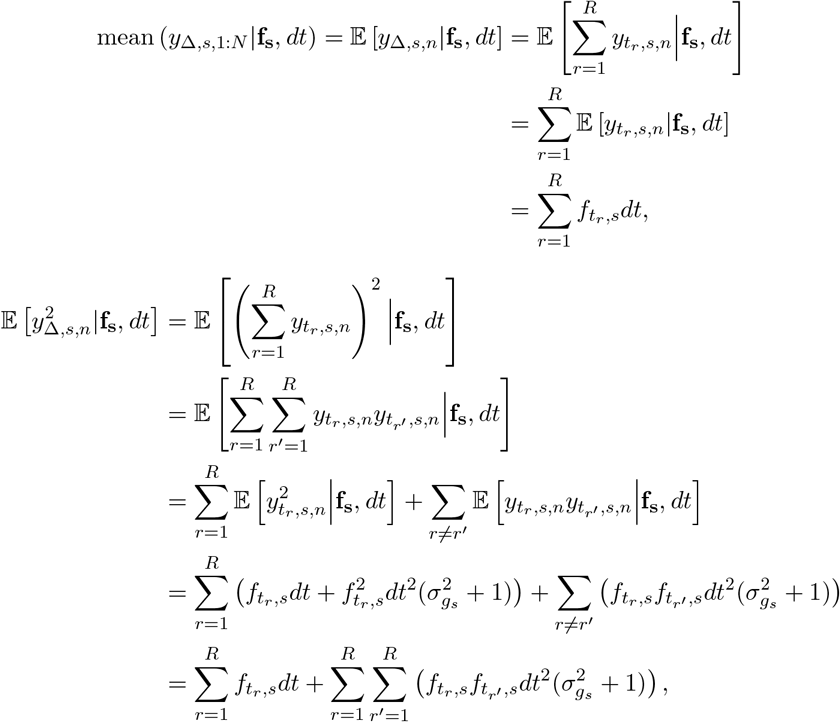

and therefore, the variance becomes:

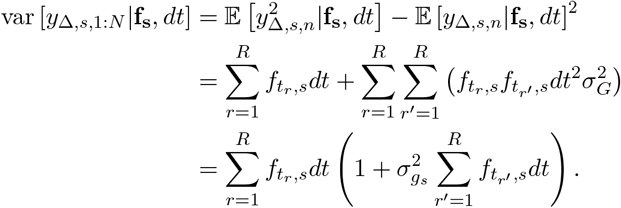

#### Appendix 3.2.2 Log-likelihood derivation

We calculate the test log-likelihood of this model using a Monte Carlo integration procedure [6]:

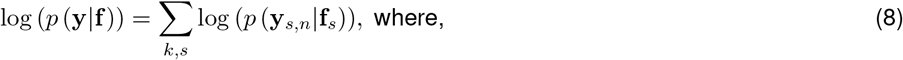

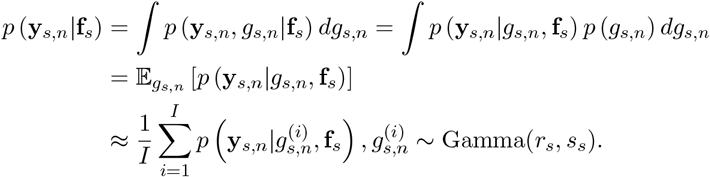

Defining:

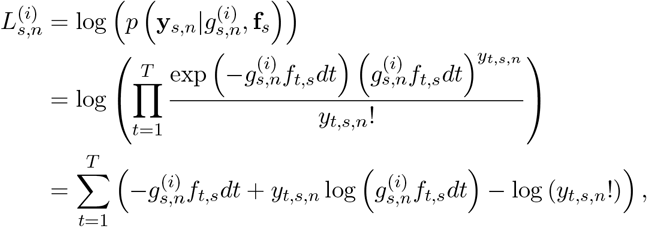

we get:

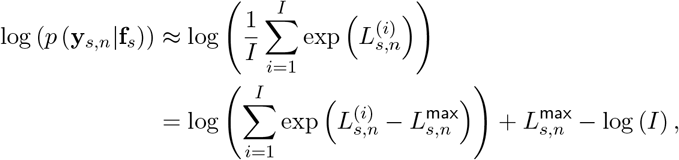

where 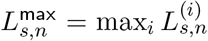. We set *I* = 1000 and use this formulation to derive the log-likelihood of the spiking observations in the *n*^th^ test trial of the *s*^th^ stimulus, and finally calculate the joint log-likelihood of all test data following Equation 8.

---

**Figure S1.**
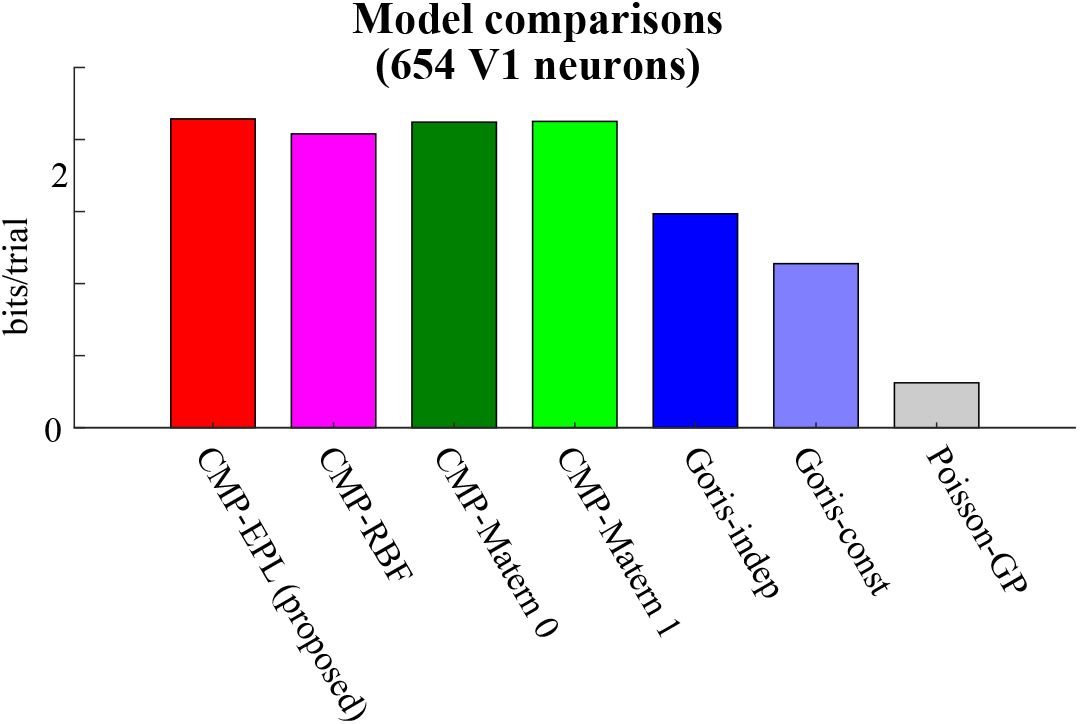
Comparison of the average test data log-likelihood improvement of each model relative to the baseline Poisson model, across 654 V1 neurons and all 72 grating orientations in recordings from the macaque primary visual cortex (data from [7]). From left to right: the proposed CMP model, the CMP-RBF model, the CMP-Matérn 0 model, the CMP-Matérn 1 model, the Goris-independent gain model, the Goris-constant gain model, and the Poisson-GP model. These results demonstrate that the CMP model systematically outperforms all other models in fitting held-out data.

**Figure S2.**
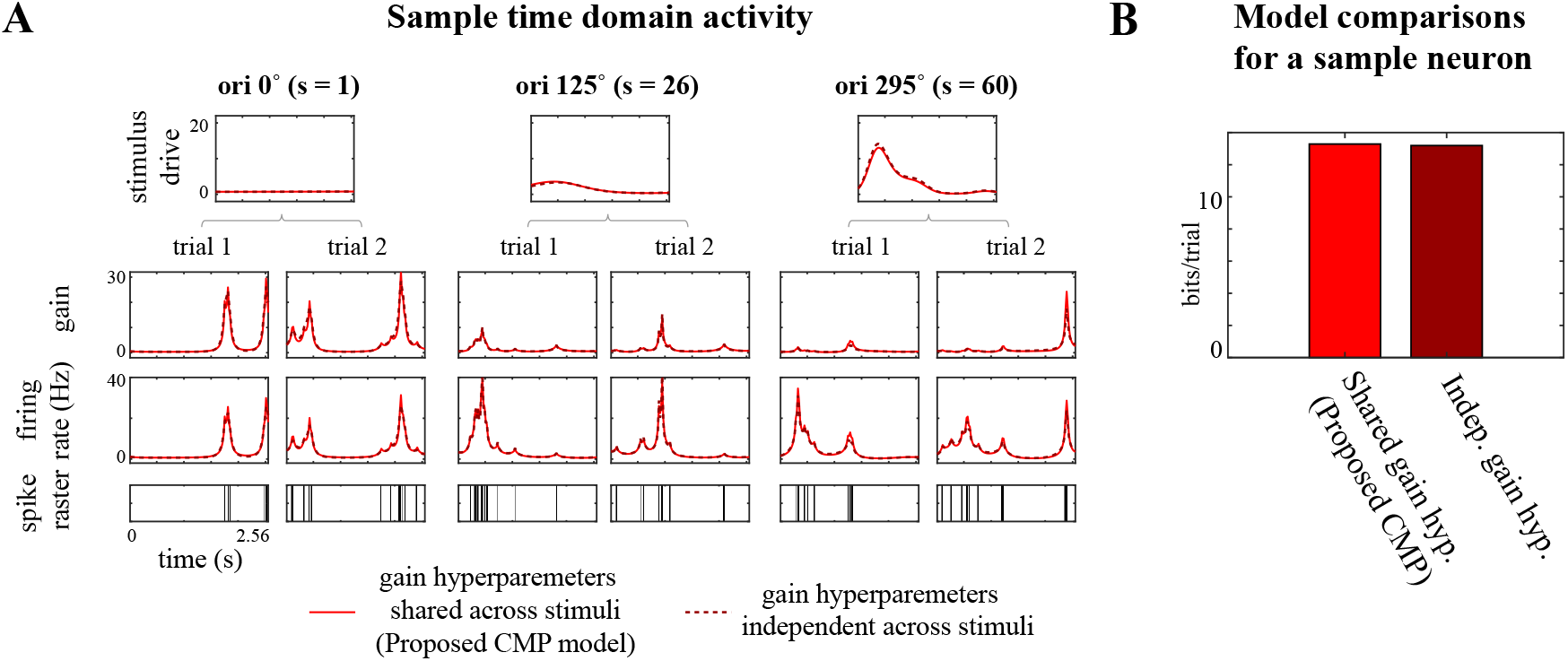
Comparison of the performance of the proposed CMP framework with an alternative model in which, instead of sharing a common set of gain hyperparameters *ρ*_*g*_, *ℓ*_*g*_, *q* across stimuli (as in the CMP framework), each stimulus condition has its own independent gain hyperparameters 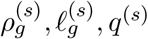. Results are shown for a sample neuron from the macaque primary visual cortex dataset (data from [7]). **(A)** Comparison of the inferred time-domain activity. From top to bottom: the inferred stimulus-driven components at three selected orientation settings (0^◦^, 125^◦^, 295^◦^); inferred gain processes for two trials per orientation; inferred firing rates from the proposed CMP model (solid lines) and the alternative model with independent gain hyperparameters per stimulus (dashed lines); and the observed spike raster. **(B)** Comparison of the log-likelihood improvement relative to the baseline Poisson model for the proposed CMP model (left) versus the alternative independent-gain model (right). These results show that the performance of both models is very similar, corroborating that the proposed CMP modeling framework robustly captures trial-to-trial variability across different stimuli using a shared set of hyperparameters.

**Figure S3.**
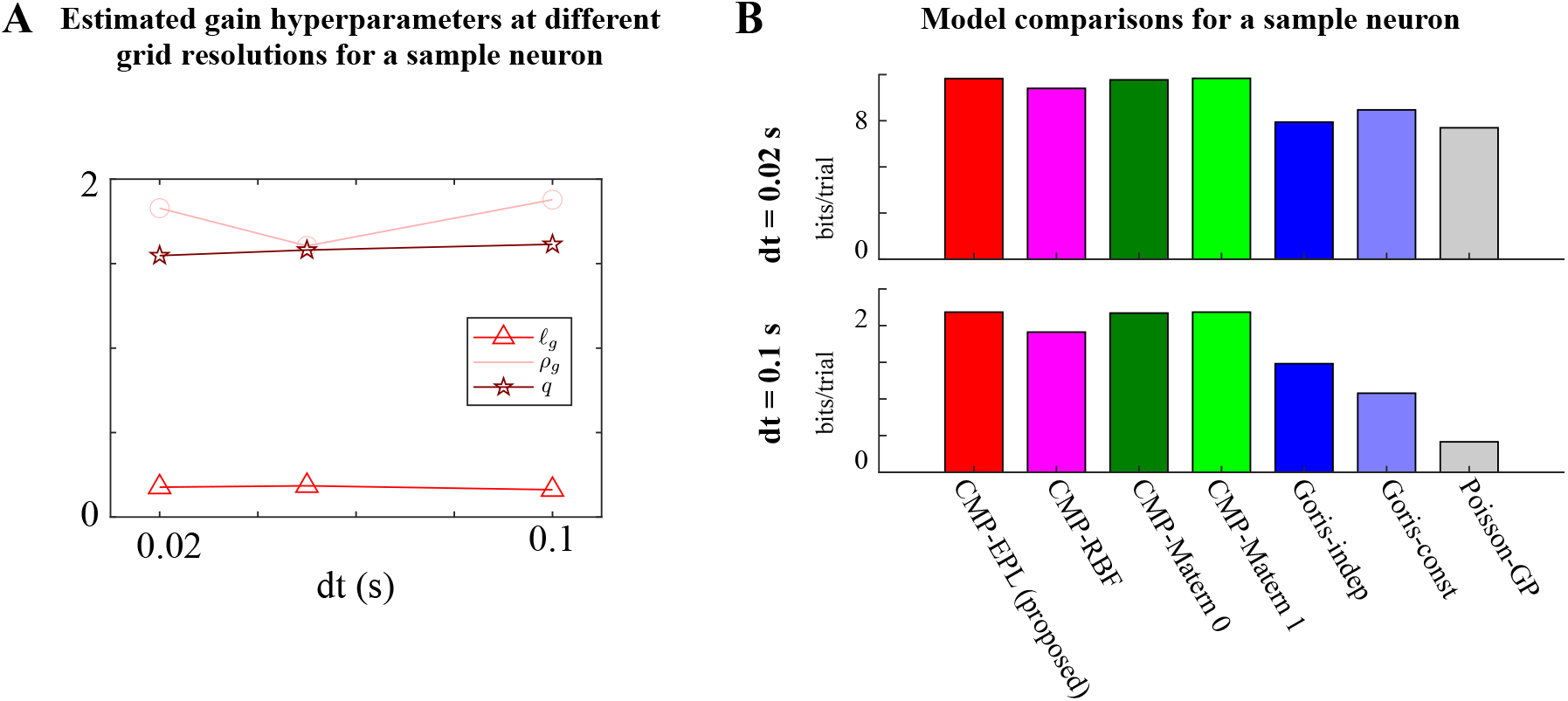
Comparison of CMP model inference at different grid sizes *dt* for a sample neuron from the macaque primary visual cortex dataset (data from [7]). **(A)** Comparison of the inferred gain hyperparameters *ℓ*_*g*_, *ρ*_*g*_, *q* across three sufficiently fine grid resolutions: *dt* = 0.02 s, *dt* = 0.05 s, and *dt* = 0.1 s. These results indicate that the inference procedure is robust to moderate changes in grid size once the temporal discretization is fine enough to approximate the underlying continuous-time model. **(B)** Comparison of the test data log-likelihood gain relative to the baseline Poisson model at two different grid resolutions (*dt* = 0.02 s and *dt* = 0.1 s). From left to right: the proposed CMP model, CMP-RBF model, CMP-Matérn 0 model, CMP-Matérn 1 model, Goris-independent gain model, Goris-constant gain model, and the Poisson-GP model. These results show that the CMP consistently outperforms alternative models over this range.

**Figure S4.**
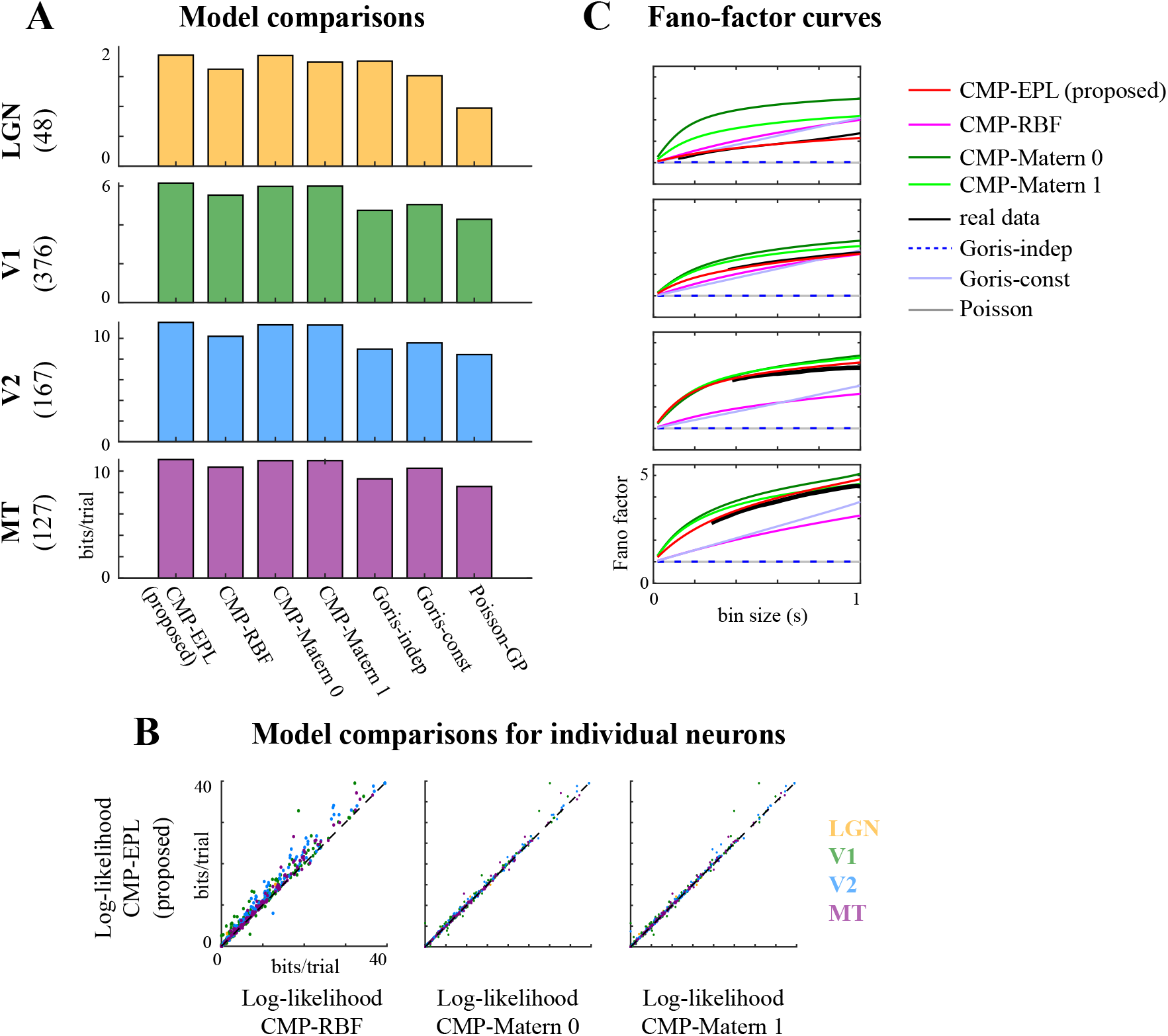
Performance comparisons of different models on neuronal population data recorded from various areas along the visual hierarchy: the lateral geniculate nucleus (LGN), V1, V2, and MT, in response to drifting sinusoidal gratings of the preferred size and speed, varying either in spatial frequency (12 spatial frequencies, ranging from 0 to 10 cycles/deg) or in drift direction (16 equally spaced directions) (data from [5]). **(A)** Log-likelihood improvement of each model relative to the baseline Poisson model, averaged across held-out data for each population. Rows from top to bottom: LGN, V1, V2, and MT. The number of neurons in each population is indicated in parentheses. From left to right: proposed CMP model, CMP-RBF model, CMP-Matérn 0 model, CMP-Matérn 1 model, Goris-independent gain model, Goris-constant gain model, and the Poisson-GP model. **(B)** Comparison of the log-likelihood of the proposed CMP model with the CMP-RBF model (left), the CMP-Matérn 0 model (middle), and the CMP-Matérn 1 model (right) for individual neurons. Colors indicate brain areas: LGN (yellow), V1 (green), V2 (blue), and MT (purple). **(C)** Comparison of the inferred Fano factor as a function of bin size, averaged across each population (rows from top to bottom: LGN, V1, V2, and MT). Curves show the real data (black), the proposed CMP model (red), CMP-RBF model (magenta), CMP-Matérn 0 model (dark green), CMP-Matérn 1 model (light green), Goris-independent gain model (dark blue dashed lines), Goris-constant gain model (light blue), and the baseline Poisson model (gray).

**Figure S5.**
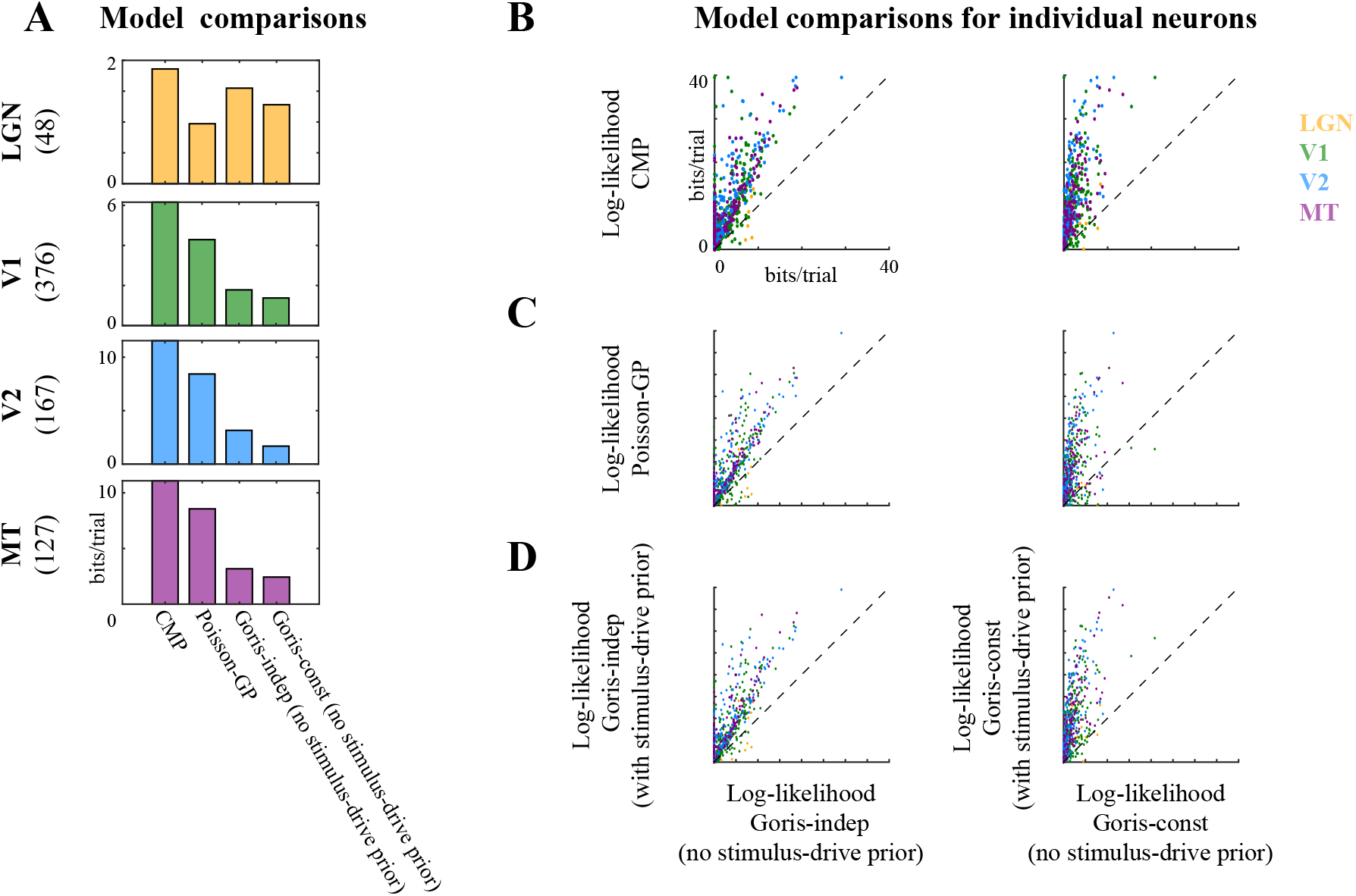
Comparison of the performance of the Goris model with and without a smoothness prior on the stimulus drive (data from [5]). **(A)** Log-likelihood improvement of each model relative to the baseline Poisson model, averaged across held-out data for each population. Rows from top to bottom: LGN, V1, V2, and MT, with the number of neurons in each population indicated in parentheses. From left to right: the proposed CMP model, the Poisson-GP model, the Goris-independent gain model without a GP prior on the stimulus drive, and the Goris-constant gain model without a GP prior on the stimulus drive. **(B)** Comparison of the log-likelihood of the proposed CMP model versus the Goris-independent gain model without a GP prior on the stimulus drive (left), and versus the Goris-constant gain model without a GP prior on the stimulus drive (right), for individual neurons. **(C)** Comparison of the log-likelihood of the Poisson-GP model versus the Goris-independent gain model without a GP prior (left), and versus the Goris-constant gain model without a GP prior (right), for individual neurons. **(D)** Comparison of the log-likelihood of the Goris-independent gain model with a GP prior on the stimulus drive versus without a GP prior (left), and the Goris-constant gain model with a GP prior versus without a GP prior (right), for individual neurons. Colors indicate the areas: LGN (yellow), V1 (green), V2 (blue), and MT (purple). These results show that including a smoothness prior on the stimulus drive generally improves the model’s ability to fit held-out data, though the proposed CMP framework still achieves systematically higher performance across all areas. This highlights the importance of jointly modeling smooth stimulus tuning and structured gain fluctuations for accurately capturing neural variability.

## References

[1] R L T Goris, J A Movshon, and E P Simoncelli. Partitioning neuronal variability. Nature Neuroscience, 17(6):858–865, 2014.

[2] George J. Tomko and Donald R. Crapper. Neuronal variability: non-stationary responses to identical visual stimuli. Brain Research, 79(3):405–418, 1974. ISSN 0006-8993. doi: 10.1016/0006-8993(74)90438-7. URL https://www.sciencedirect.com/science/article/pii/0006899374904387.

[3] G.H. Henry, P.O. Bishop, R.M. Tupper, and B. Dreher. Orientation specificity and response variability of cells in the striate cortex. Vision Research, 13(9):1771–1779, 1973. ISSN 0042-6989. doi: 10.1016/0042-6989(73)90094-1. URL https://www.sciencedirect.com/science/article/pii/0042698973900941.

[4] M. Carandini. Amplification of trial-to-trial response variability by neurons in visual cortex. PLoS biology, 2(9):e264, 2004.

[5] Ilan Dinstein, David J. Heeger, and Marlene Behrmann. Neural variability: friend or foe? Trends in Cognitive Sciences, 19(6):322–328, 2015. ISSN 1364-6613. doi: 10.1016/j.tics.2015.04.005. URL https://www.sciencedirect.com/science/article/pii/S1364661315000911.

[6] C Allen and C F Stevens. An evaluation of causes for unreliability of synaptic transmission. Proceedings of the National Academy of Sciences, 91(22):10380–10383, 1994. doi: 10.1073/pnas.91.22.10380. URL https://www.pnas.org/doi/abs/10.1073/pnas.91.22.10380.

[7] Michael N. Shadlen and William T. Newsome. The variable discharge of cortical neurons: Implications for connectivity, computation, and information coding. Journal of Neuroscience, 18 (10):3870–3896, 1998. ISSN 0270-6474. doi: 10.1523/JNEUROSCI.18-10-03870.1998. URL https://www.jneurosci.org/content/18/10/3870.

[8] David M. Schneeweis and Julie L. Schnapf. The photovoltage of macaque cone photoreceptors: Adaptation, noise, and kinetics. Journal of Neuroscience, 19(4):1203–1216, 1999. ISSN 0270-6474. doi: 10.1523/JNEUROSCI.19-04-01203.1999. URL https://www.jneurosci.org/content/19/4/1203.

[9] C. van Vreeswijk and H. Sompolinsky. Chaos in neuronal networks with balanced excitatory and inhibitory activity. Science, 274(5293):1724–1726, 1996. doi: 10.1126/science.274.5293.1724. URL https://www.science.org/doi/abs/10.1126/science.274.5293.1724.

[10] Colin W.G. Clifford, Michael A. Webster, Garrett B. Stanley, Alan A. Stocker, Adam Kohn, Tatyana O. Sharpee, and Odelia Schwartz. Visual adaptation: Neural, psychological and computational aspects. Vision Research, 47(25):3125–3131, 2007. ISSN 0042-6989. doi: 10.1016/j.visres.2007.08.023. URL https://www.sciencedirect.com/science/article/pii/S0042698907003756.

[11] Daniel E. Feldman. Synaptic mechanisms for plasticity in neocortex. Annual Review of Neuroscience, 32(1):33–55, 2009. doi: 10.1146/annurev.neuro.051508.135516. URL https://doi.org/10.1146/annurev.neuro.051508.135516. PMID: 19400721.

[12] Donald H. Perkel, George L. Gerstein, and George P. Moore. Neuronal spike trains and stochastic point processes: I. the single spike train. Biophysical Journal, 7(4):391–418, 1967. ISSN 0006-3495. doi: 10.1016/S0006-3495(67)86596-2. URL https://www.sciencedirect.com/science/article/pii/S0006349567865962.

[13] Wilson Truccolo, Uri T. Eden, Matthew R. Fellows, John P. Donoghue, and Emery N. Brown. A point process framework for relating neural spiking activity to spiking history, neural ensemble, and extrinsic covariate effects. Journal of Neurophysiology, 93(2):1074–1089, 2005. doi: 10.1152/jn.00697.2004. URL https://doi.org/10.1152/jn.00697.2004. PMID: 15356183.

[14] Liam Paninski, Jonathan Pillow, and Jeremy Lewi. Statistical models for neural encoding, decoding, and optimal stimulus design. In Paul Cisek, Trevor Drew, and John F. Kalaska, editors, Computational Neuroscience: Theoretical Insights into Brain Function, volume 165 of Progress in Brain Research, pages 493–507. Elsevier, 2007. doi: 10.1016/S0079-6123(06)65031-0. URL https://www.sciencedirect.com/science/article/pii/S0079612306650310.

[15] J. W. Pillow. Time-rescaling methods for the estimation and assessment of non-poisson neural encoding models. In Y. Bengio, D. Schuurmans, J. Lafferty, C. K. I. Williams, and A. Culotta, editors, Advances in Neural Information Processing Systems 22, pages 1473–1481. MIT Press, 2009.

[16] L. Paninski, E. N. Brown, S. Iyengar, R. E. Kass, Liam Paninski, Emery N Brown, Satish Iyengar, and Robert E Kass. Statistical models of spike trains. Stochastic methods in neuroscience, 24: 278–303, 2009.

[17] Alison I. Weber and Jonathan W. Pillow. Capturing the dynamical repertoire of single neurons with generalized linear models. Neural Computation, 29(12):3260–3289, 2017. doi: 10.1162/neco_a_01021. URL https://doi.org/10.1162/neco_a_01021.

[18] Henry C. Tuckwell. Introduction to Theoretical Neurobiology. Cambridge Studies in Mathematical Biology. Cambridge University Press, 1988.

[19] Fred Rieke, Davd Warland, Rob de Ruyter van Steveninck, and William Bialek. Spikes: exploring the neural code. MIT Press, Cambridge, MA, USA, 1999. ISBN 0262181746.

[20] N. V. Swindale and D. E. Mitchell. Comparison of receptive field properties of neurons in area 17 of normal and bilaterally amblyopic cats. Experimental Brain Research, 99(3):399–410, Jan 1994. ISSN 1432-1106. doi: 10.1007/BF00228976. URL https://doi.org/10.1007/BF00228976.

[21] Roland Baddeley, L. F. Abbott, Michael C. A. Booth, Frank Sengpiel, Tobe Freeman, Edward A. Wakeman, and Edmund T. Rolls. Responses of neurons in primary and inferior temporal visual cortices to natural scenes. Proceedings of the Royal Society of London. Series B: Biological Sciences, 264(1389):1775–1783, 1997. doi: 10.1098/rspb.1997.0246. URL https://royalsocietypublishing.org/doi/abs/10.1098/rspb.1997.0246.

[22] P. Lánsky and J. Vaillant. Stochastic model of the overdispersion in the place cell discharge. Biosystems, 58(1):27–32, 2000.

[23] Jonathan Pillow and James Scott. Fully bayesian inference for neural models with negative-binomial spiking. In F. Pereira, C.J. Burges, L. Bottou, and K.Q. Weinberger, editors, Advances in Neural Information Processing Systems, volume 25. Curran Associates, Inc., 2012. URL https://proceedings.neurips.cc/paper_files/paper/2012/file/b55ec28c52d5f6205684a473a2193564-Paper.pdf.

[24] Yuanjun Gao, Lars Busing, Krishna V Shenoy, and John P Cunningham. High-dimensional neural spike train analysis with generalized count linear dynamical systems. In C. Cortes, N. Lawrence, D. Lee, M. Sugiyama, and R. Garnett, editors, Advances in Neural Information Processing Systems, volume 28. Curran Associates, Inc., 2015. URL https://proceedings.neurips.cc/paper_files/paper/2015/file/9996535e07258a7bbfd8b132435c5962-Paper.pdf.

[25] Ian H. Stevenson. Flexible models for spike count data with both over- and under-dispersion. Journal of Computational Neuroscience, 41(1):29–43, Aug 2016. ISSN 1573-6873. doi: 10.1007/s10827-016-0603-y. URL https://doi.org/10.1007/s10827-016-0603-y.

[26] Adam S. Charles, Mijung Park, J. Patrick Weller, Gregory D. Horwitz, and Jonathan W. Pillow. Dethroning the Fano Factor: A Flexible, Model-Based Approach to Partitioning Neural Variability. Neural Computation, 30(4):1012–1045, 04 2018. ISSN 0899-7667. doi: 10.1162/neco_a_01062. URL https://doi.org/10.1162/neco_a_01062.

[27] R. Barbieri, M. C. Quirk, L. M. Frank, M. A. Wilson, and E. N. Brown. Construction and analysis of non-poisson stimulus-response models of neural spiking activity. Journal of Neuroscience Methods, 105(1):25–37, 2001.

[28] Qi She, Xiaoli Wu, Beth Jelfs, Adam S. Charles, and Rosa H. M. Chan. An efficient and flexible spike train model via empirical bayes. IEEE Transactions on Signal Processing, 69:3236–3251, 2021. doi: 10.1109/TSP.2021.3076885.

[29] Cina Aghamohammadi, Chandramouli Chandrasekaran, and Tatiana A. Engel. A doubly stochastic renewal framework for partitioning spiking variability. bioRxiv, 2024.

[30] A Charles and J Pillow. Additive continuous-time joint partitioning of neural variability. In 2018 Conference on Cognitive Computational Neuroscience. 10.32470/CCN, 2018.

[31] Robbe L. T. Goris, Corey M. Ziemba, J. Anthony Movshon, and Eero P. Simoncelli. Slow gain fluctuations limit benefits of temporal integration in visual cortex. Journal of Vision, 18(8):8–8, 08 2018. ISSN 1534-7362. doi: 10.1167/18.8.8. URL https://doi.org/10.1167/18.8.8.

[32] Olivier J Hénaff, Zoe M Boundy-Singer, Kristof Meding, Corey M Ziemba, and Robbe L T Goris. Representation of visual uncertainty through neural gain variability. Nat. Commun., 11(1):2513, May 2020.

[33] Carl Edward Rasmussen and Christopher K. I. Williams. Gaussian processes for machine learning. Adaptive computation and machine learning. MIT Press, 2006. ISBN 026218253X.

[34] Yuan Zhao and Il Memming Park. Variational Latent Gaussian Process for Recovering Single-Trial Dynamics from Population Spike Trains. Neural Computation, 29(5):1293–1316, 05 2017. ISSN 0899-7667. doi: 10.1162/NECO_a_00953. URL https://doi.org/10.1162/NECO_a_00953.

[35] Stephen Keeley, Mikio Aoi, Yiyi Yu, Spencer Smith, and Jonathan W Pillow. Identifying signal and noise structure in neural population activity with gaussian process factor models. In H. Larochelle, M. Ranzato, R. Hadsell, M.F. Balcan, and H. Lin, editors, Advances in Neural Information Processing Systems, volume 33, pages 13795–13805. Curran Associates, Inc., 2020.

[36] A B A Graf, A Kohn, M Jazayeri, and J A Movshon. Decoding the activity of neuronal populations in macaque primary visual cortex. Nature Neuroscience, 14(2):239–245, Feb 2011. doi: 10.1038/nn.2733. URL https://doi.org/10.1038/nn.2733.

[37] D. R. Cox. Some statistical methods connected with series of events. Journal of the Royal Statistical Society: Series B (Methodological), 17(2):129–157, 1955. doi: 10.1111/j.2517-6161.1955.tb00188.x. URL https://rss.onlinelibrary.wiley.com/doi/abs/10.1111/j.2517-6161.1955.tb00188.x.

[38] Michael I Jordan, Zoubin Ghahramani, Tommi S Jaakkola, and Lawrence K Saul. An introduction to variational methods for graphical models. Machine Learning, 37(2):183–233, 1999. doi: 10.1023/A:1007665907178.

[39] Mark M Churchland, Byron M Yu, John P Cunningham, Leo P Sugrue, Marlene R Cohen, Greg S Corrado, William T Newsome, Andrew M Clark, Paymon Hosseini, Benjamin B Scott, David C Bradley, Matthew A Smith, Adam Kohn, J Anthony Movshon, Katherine M Armstrong, Tirin Moore, Steve W Chang, Lawrence H Snyder, Stephen G Lisberger, Nicholas J Priebe, Ian M Finn, David Ferster, Stephen I Ryu, Gopal Santhanam, Maneesh Sahani, and Krishna V Shenoy. Stimulus onset quenches neural variability: a widespread cortical phenomenon. Nat. Neurosci., 13(3):369–378, March 2010.

[40] Robbe L T Goris, Ruben Coen-Cagli, Kenneth D Miller, Nicholas J Priebe, and Máté Lengyel. Response sub-additivity and variability quenching in visual cortex. Nat. Rev. Neurosci., 25(4): 237–252, April 2024.

[41] Oleg I Rumyantsev, Jérôme A Lecoq, Oscar Hernandez, Yanping Zhang, Joan Savall, Radosław Chrapkiewicz, Jane Li, Hongkui Zeng, Surya Ganguli, and Mark J Schnitzer. Fundamental bounds on the fidelity of sensory cortical coding. Nature, 580(7801):100—105, April 2020. ISSN 0028-0836. URL 10.1038/s41586-020-2130-2.

[42] Ramon Bartolo, Richard C. Saunders, Andrew R. Mitz, and Bruno B. Averbeck. Information-limiting correlations in large neural populations. Journal of Neuroscience, 40(8):1668–1678, 2020. ISSN 0270-6474. doi: 10.1523/JNEUROSCI.2072-19.2019. URL https://www.jneurosci.org/content/40/8/1668.

[43] Anuththara Rupasinghe, Nikolas Francis, Ji Liu, Zac Bowen, Patrick O Kanold, and Behtash Babadi. Direct extraction of signal and noise correlations from two-photon calcium imaging of ensemble neuronal activity. eLife, 10:e68046, jun 2021. ISSN 2050-084X. doi: 10.7554/eLife.68046. URL https://doi.org/10.7554/eLife.68046.

[44] James J. Jun, Nick Steinmetz, Joshua H. Siegle, Daniel J. Denman, Marius Bauza, Brian Barbarits, Albert K. Lee, Costas Anastassiou, Alexandru Andrei, Cagatay Aydin, Mladen Barbic, Tim Blanche, Vincent Bonin, Joao Couto, Barundeb Dutta, Sergey Gratiy, Diego Gutnisky, Michael Häusser, Bill Karsh, and Timothy D. Harris. Fully integrated silicon probes for high-density recording of neural activity. Nature, 551:232–236, 11 2017. doi: 10.1038/nature24636.

[45] Jiangang Du, Timothy J. Blanche, Reid R. Harrison, Henry A. Lester, and Sotiris C. Masmanidis. Multiplexed, high density electrophysiology with nanofabricated neural probes. PLOS ONE, 6(10): 1–11, 10 2011. doi: 10.1371/journal.pone.0026204.

[46] David Liu and Máté Lengyel. A universal probabilistic spike count model reveals ongoing modulation of neural variability. Advances in Neural Information Processing Systems, 34, 2021.

[47] Sacha Sokoloski, Amir Aschner, and Ruben Coen-Cagli. Modelling the neural code in large populations of correlated neurons. eLife, 10:e64615, oct 2021. ISSN 2050-084X. doi: 10.7554/eLife.64615. URL https://doi.org/10.7554/eLife.64615.

[48] A. Aldo Faisal, Luc P. J. Selen, and Daniel M. Wolpert. Noise in the nervous system. Nature Reviews Neuroscience, 9(4):292–303, Apr 2008. ISSN 1471-0048. doi: 10.1038/nrn2258. URL https://doi.org/10.1038/nrn2258.

[49] Eero P Simoncelli and Bruno A Olshausen. Natural image statistics and neural representation. Annual review of neuroscience, 24(1):1193–1216, 2001.

[50] Gergő Orbán, Pietro Berkes, József Fiser, and Máté Lengyel. Neural variability and sampling-based probabilistic representations in the visual cortex. Neuron, 92(2):530–543, 2016. ISSN 0896-6273. doi: 10.1016/j.neuron.2016.09.038. URL https://www.sciencedirect.com/science/article/pii/S0896627316306390.

[51] Mark M Churchland, Byron M Yu, Stephen I Ryu, Gopal Santhanam, and Krishna V Shenoy. Neural variability in premotor cortex provides a signature of motor preparation. J Neurosci, 26(14): 3697–3712, April 2006.

[52] C. W. Gardiner. Handbook of stochastic methods for physics, chemistry and the natural sciences, volume 13 of Springer Series in Synergetics. Springer-Verlag, Berlin, third edition, 2004. ISBN 3–540-20882-8.

[53] G. E. Uhlenbeck and L. S. Ornstein. On the theory of the brownian motion. Phys. Rev., 36:823–841, Sep 1930. doi: 10.1103/PhysRev.36.823. URL https://link.aps.org/doi/10.1103/PhysRev.36.823.

[54] Benoit B. Mandelbrot and John W. Van Ness. Fractional brownian motions, fractional noises and applications. SIAM Review, 10(4):422–437, 1968. ISSN 00361445. URL http://www.jstor.org/stable/2027184.

[55] Yuhai Tu and G. Grinstein. How white noise generates power-law switching in bacterial flagellar motors. Phys. Rev. Lett., 94:208101, May 2005. doi: 10.1103/PhysRevLett.94.208101. URL https://link.aps.org/doi/10.1103/PhysRevLett.94.208101.

[56] Alberto Stefano Sassi, Mayra Garcia-Alcala, Maximino Aldana, and Yuhai Tu. Protein concentration fluctuations in the high expression regime: Taylor’s law and its mechanistic origin. Phys. Rev. X, 12:011051, Mar 2022. doi: 10.1103/PhysRevX.12.011051. URL https://link.aps.org/doi/10.1103/PhysRevX.12.011051.

[57] M J Berry, 2nd and M Meister. Refractoriness and neural precision. J. Neurosci., 18(6):2200–2211, March 1998.

[58] Gaby Maimon and John A Assad. Beyond poisson: increased spike-time regularity across primate parietal cortex. Neuron, 62(3):426–440, May 2009.

[59] Ulisse Ferrari, Stéphane Deny, Olivier Marre, and Thierry Mora. A simple model for low variability in neural spike trains. Neural Computation, 30(11):3009–3036, 11 2018. ISSN 0899-7667. doi: 10.1162/neco_a_01125. URL https://doi.org/10.1162/neco_a_01125.

[60] Marlene R. Cohen and John H. R. Maunsell. Attention improves performance primarily by reducing interneuronal correlations. Nature Neuroscience, 12(12):1594–1600, Dec 2009. ISSN 1546-1726. doi: 10.1038/nn.2439. URL https://doi.org/10.1038/nn.2439.

[61] Matthew J. McGinley, Martin Vinck, Jacob Reimer, Renata Batista-Brito, Edward Zagha, Cathryn R. Cadwell, Andreas S. Tolias, Jessica A. Cardin, and David A. McCormick. Waking state: Rapid variations modulate neural and behavioral responses. Neuron, 87(6):1143–1161, 2015. ISSN 0896-6273. doi: 10.1016/j.neuron.2015.09.012. URL https://www.sciencedirect.com/science/article/pii/S0896627315007692.

[62] Carsen Stringer, Marius Pachitariu, Nicholas Steinmetz, Charu Bai Reddy, Matteo Carandini, and Kenneth D. Harris. Spontaneous behaviors drive multidimensional, brainwide activity. Science, 364(6437):eaav7893, 2019. doi: 10.1126/science.aav7893. URL https://www.science.org/doi/abs/10.1126/science.aav7893.

[63] Joseph M. Hilbe. Negative Binomial Regression. Cambridge University Press, 2 edition, 2011.

[64] A. Colin Cameron and Pravin K. Trivedi. Regression Analysis of Count Data. Econometric Society Monographs. Cambridge University Press, 2 edition, 2013.

[65] J k. Lindsey. Modelling Frequency and Count Data. Oxford University Press, 04 1995. ISBN 9780198523314. doi: 10.1093/oso/9780198523314.001.0001. URL https://doi.org/10.1093/oso/9780198523314.001.0001.

[66] Anuththara Rupasinghe. Continous partitioning of neuronal variability: Matlab codes. https://github.com/Anuththara-Rupasinghe/CMP, 2024.

[67] Mikio C. Aoi and Jonathan W. Pillow. Scalable bayesian inference for high-dimensional neural receptive fields. bioRxiv, 2017. doi: 10.1101/212217.

[68] Yung Liang Tong. The multivariate normal distribution. Springer Science & Business Media, 2012.

[69] Surya T. Tokdar and Robert E. Kass. Importance sampling: a review. WIREs Computational Statistics, 2(1):54–60, 2010. doi: 10.1002/wics.56. URL https://wires.onlinelibrary.wiley.com/doi/abs/10.1002/wics.56.

[70] Persi Diaconis and Donald Ylvisaker. Conjugate Priors for Exponential Families. The Annals of Statistics, 7(2):269 –281, 1979. doi: 10.1214/aos/1176344611. URL https://doi.org/10.1214/aos/1176344611.

## References

[1] Carl Edward Rasmussen and Christopher K. I. Williams. Gaussian processes for machine learning. Adaptive computation and machine learning. MIT Press, 2006. ISBN 026218253X.

[2] I. J. Schoenberg. Metric spaces and positive definite functions. Transactions of the American Mathematical Society, 44(3):522–536, 1938. ISSN 00029947, 10886850. URL http://www.jstor.org/stable/1989894.

[3] G.B. Folland. Real Analysis: Modern Techniques and Their Applications. Pure and Applied Mathematics: A Wiley Series of Texts, Monographs and Tracts. Wiley, 2013. ISBN 9781118626399. URL https://books.google.com/books?id=wI4fAwAAQBAJ.

[4] George Casella and Roger L. Berger. Statistical Inference. Duxbury Press, Pacific Grove, CA, 2nd edition, 2002.

[5] R L T Goris, J A Movshon, and E P Simoncelli. Partitioning neuronal variability. Nature Neuroscience, 17(6):858–865, 2014.

[6] Christian P Robert, George Casella, and George Casella. Monte Carlo statistical methods, volume 2. Springer, 1999.

[7] A B A Graf, A Kohn, M Jazayeri, and J A Movshon. Decoding the activity of neuronal populations in macaque primary visual cortex. Nature Neuroscience, 14(2):239–245, Feb 2011. doi: 10.1038/nn.2733. URL https://doi.org/10.1038/nn.2733.

