## Supplementary material for "Continuous partitioning of neuronal variability": SI Appendix

### Continuous partitioning of neuronal variability Supporting Information

#### 1 Proof of the validity of the EPL covariance function

In this section, we provide the formal proof that the proposed Exponetiated Power Law (EPL) kernel is a valid covariance function.

**Theorem 1** (Validity of the EPL Kernel). *The Exponetiated Power Law (EPL) kernel given by:*

$$K_g(t_i, t_j) = \rho_g \exp\left(-\frac{1}{2}\left|\frac{t_i - t_j}{\ell_g}\right|^q\right), \tag{1}$$

*where $\rho_g > 0$ is the marginal variance, $\ell_g > 0$ is the length scale, and $0 < q \leq 2$ is the power law exponent, is a valid covariance function.*

*Proof.* We begin by recalling the definition of a valid covariance function [1].

**Definition 1** (Covariance function). *A (real-valued) function*

$$K : \mathbb{R} \times \mathbb{R} \ \rightarrow \ \mathbb{R},$$

*is called a* covariance kernel *(or* positive semidefinite kernel*) if, for every finite set of points $t_1, \ldots, t_n \in \mathbb{R}$ and all real weights $a_1, \ldots, a_n \in \mathbb{R}$, the following Gram sum is non-negative:*

$$\sum_{i=1}^{n}\sum_{j=1}^{n} a_i \, a_j \, K(t_i, t_j) \ \geq \ 0.$$

We now invoke the following result from the Schoenberg's theorem [2].

**Lemma 1** (Schoenberg (1938)). *The function $\exp(-|x|^q)$ is positive definite on $\mathbb{R}$ for all $0 < q \leq 2$.*

*Proof of Lemma 1.* The proof is provided in [2]. □

From Lemma 1 and Definition 1, it follows that for any finite set $\tilde{t}_1, \ldots, \tilde{t}_n \in \mathbb{R}$ and weights $a_1, \ldots, a_n \in \mathbb{R}$,

$$\sum_{i=1}^{n} \sum_{j=1}^{n} a_i \, a_j \, \exp\left(-|\tilde{t}_i - \tilde{t}_j|^q\right) \; \geq \; 0,$$

if $0 < q \leq 2$.

Next, consider the change of variables $t_i = 2^{1/q} \ell_g \cdot \tilde{t}_i$, which implies:

$$\left|\tilde{t}_i - \tilde{t}_j\right|^q = \left| \frac{t_i}{(2)^{1/q}\ell_g} - \frac{t_j}{(2)^{1/q}\ell_g} \right|^q = \frac{1}{2} \left| \frac{t_i - t_j}{\ell_g} \right|^q$$

Substituting this into the above expression, we obtain:

$$\sum_{i=1}^{n} \sum_{j=1}^{n} a_i a_j \exp\left( -\frac{1}{2} \left| \frac{t_i - t_j}{\ell_g} \right|^q \right) \geq 0.$$

Finally, since $\rho_g > 0$, multiplying by $\rho_g$ preserves non-negativity:

$$\sum_{i=1}^{n} \sum_{j=1}^{n} a_i a_j \rho_g \exp\left( -\frac{1}{2} \left| \frac{t_i - t_j}{\ell_g} \right|^q \right) \geq 0.$$

Hence, for any finite set $t_1, \ldots, t_n \in \mathbb{R}$ and any real weights $a_1, \ldots, a_n \in \mathbb{R}$, we have:

$$\sum_{i=1}^{n} \sum_{j=1}^{n} a_i \, a_j \, K_g(t_i, t_j) \; \geq \; 0,$$

which establishes that the EPL kernel in Equation 1 is positive semidefinite, and thus a valid covariance function. □

#### 2 Partitioning variability with CMP

In this section, we provide the detailed derivations of the mean–variance relationship under the CMP model. Under the CMP model, spike train variability can be partitioned into a Poisson component, which is equal to the mean, and a modulatory component, which controls the degree of overdispersion. The continuous-time formulation of the CMP model allows us to compute this partitioning for arbitrary time intervals ($\Delta$), and thus to analytically characterize how overdispersion grows with time bin size.

Recall the CMP forward model:

$$\begin{aligned} y(t) \mid s &\sim \text{PoissProc}\big(\lambda(t)\big), \\ \lambda(t) &= f_s(t) \cdot g(t), \\ \tilde{g}(t) &:= \log g(t) \sim \mathcal{GP}(0, K_g), \\ K_g(t_1, t_2) &= \rho_g \exp\left( -\frac{1}{2} \left| \frac{t_1 - t_2}{\ell_g} \right|^q \right). \end{aligned}$$

Under this model, the mean spike count in a time bin of size $\Delta$, conditioned on the stimulus $s$, can be derived as:

$$\text{mean}[y_\Delta \mid s] = \mathbb{E}\left[y_\Delta \mid s\right]$$

$$
\begin{aligned}
&= \mathbb{E}\left[\int_0^{\Delta} y(t)dt \mid s\right] \\
&\overset{(a)}{=} \int_0^{\Delta} \mathbb{E}\left[y(t) \mid s\right] dt \\
&\overset{(b)}{=} \int_0^{\Delta} \mathbb{E}\left[\lambda(t) \mid s\right] dt \\
&= \int_0^{\Delta} \mathbb{E}\left[f_s(t) \cdot g(t) \mid s\right] dt \\
&= \int_0^{\Delta} f_s(t)\mathbb{E}\left[g(t)\right] dt \\
&= \int_0^{\Delta} f_s(t)\mathbb{E}\left[\exp\left(\tilde{g}(t)\right)\right] dt \\
&\overset{(c)}{=} \int_0^{\Delta} f_s(t)\exp\left(\rho_g/2\right) dt \\
&= \exp\left(\rho_g/2\right)\int_0^{\Delta} f_s(t)dt,
\end{aligned} \tag{2}
$$

where $(a)$ follows from Fubini's Theorem [3], $(b)$ uses the fact that the mean of a Poisson process equals its rate function [3], and $(c)$ applies the moment-generating function of a Gaussian (i.e., the mean of a log-normal distribution) [4].

Using similar steps, the second moment can be derived as:

$$
\begin{aligned}
\mathbb{E}\left[y_{\Delta}^2 \mid s\right] &= \mathbb{E}\left[\left(\int_0^{\Delta} y(t)dt\right)^2 \mid s\right] \\
&= \mathbb{E}\left[\iint y(t_1)y(t_2)\, dt_1 dt_2 \mid s\right] \\
&= \iint \mathbb{E}\left[y(t_1)y(t_2) \mid s\right] dt_1 dt_2 \\
&= \iint_{t_1=t_2} \mathbb{E}\left[y(t_1)y(t_2) \mid s\right] dt_1 dt_2 + \iint_{t_1\neq t_2} \mathbb{E}\left[y(t_1)y(t_2) \mid s\right] dt_1 dt_2 \\
&= \iint_{t_1=t_2} \mathbb{E}\left[y(t_1)^2 \mid s\right] dt_1 dt_2 + \iint_{t_1\neq t_2} \mathbb{E}\left[y(t_1)y(t_2) \mid s\right] dt_1 dt_2 \\
&= \iint_{t_1=t_2} \mathbb{E}\left[\lambda(t_1) + \lambda(t_1)^2 \mid s\right] dt_1 dt_2 + \iint_{t_1\neq t_2} \mathbb{E}\left[\lambda(t_1)\lambda(t_2) \mid s\right] dt_1 dt_2 \\
&= \iint_{t_1=t_2} \left(f_s(t_1)\mathbb{E}\left[g(t_1)\right] + f_s(t_1)^2\mathbb{E}\left[g(t_1)^2\right]\right) dt_1 dt_2 + \iint_{t_1\neq t_2} f_s(t_1)f_s(t_2)\mathbb{E}\left[g(t_1)g(t_2)\right] dt_1 dt_2 \\
&= \iint_{t_1=t_2} \left(f_s(t_1)\mathbb{E}\left[\exp\left(\tilde{g}(t_1)\right)\right] + f_s(t_1)^2\mathbb{E}\left[\exp\left(2\tilde{g}(t_1)\right)\right]\right) dt_1 dt_2 \\
&\qquad + \iint_{t_1\neq t_2} f_s(t_1)f_s(t_2)\mathbb{E}\left[\exp\left(\tilde{g}(t_1) + \tilde{g}(t_2)\right)\right] dt_1 dt_2 \\
&= \iint_{t_1=t_2} \left(f_s(t_1)\exp\left(\rho_g/2\right) + f_s(t_1)^2\exp\left(2\rho_g\right)\right) dt_1 dt_2
\end{aligned}
$$

$$
\begin{aligned}
&+ \iint_{t_1 \neq t_2} f_s(t_1) f_s(t_2) \exp\left(\rho_g + K_g(t_1, t_2)\right) dt_1 dt_2 \\
&= \exp\left(\rho_g/2\right) \int_0^{\Delta} f_s(t) dt + \exp\left(\rho_g\right) \iint f_s(t_1) f_s(t_2) \exp\left(K_g(t_1, t_2)\right) dt_1 dt_2. \quad (3)
\end{aligned}
$$

Thus, using Equation 2 and Equation 3 the variance can be formulated as:

$$
\begin{aligned}
\operatorname{var}[y_\Delta \mid s] &= \mathbb{E}\left[y_\Delta^2 \mid s\right] - \left(\mathbb{E}\left[y_\Delta \mid s\right]\right)^2 \\
&= \exp\left(\rho_g/2\right) \int_0^{\Delta} f_s(t) dt + \exp\left(\rho_g\right) \iint f_s(t_1) f_s(t_2) \exp\left(K_g(t_1, t_2)\right) dt_1 dt_2 \\
&\quad - \left(\exp\left(\rho_g/2\right) \int_0^{\Delta} f_s(t) dt\right) \\
&= \exp\left(\rho_g/2\right) \int_0^{\Delta} f_s(t) dt + \exp\left(\rho_g\right) \iint f_s(t_1) f_s(t_2) \left(\exp\left(K_g(t_1, t_2)\right) - 1\right) dt_1 dt_2. \quad (4)
\end{aligned}
$$

From these expressions, we can compute Fano factor as a function of bin size, which is simply the ratio of variance (Equation 4) to mean (Equation 2):

$$
\begin{aligned}
FF(\Delta) &= \frac{\operatorname{var}[y_\Delta \mid s]}{\mathbb{E}\left[y_\Delta \mid s\right]} \\
&= \frac{\exp\left(\rho_g/2\right) \int_0^{\Delta} f_s(t) dt + \exp\left(\rho_g\right) \iint f_s(t_1) f_s(t_2) \left(\exp\left(K_g(t_1, t_2)\right) - 1\right) dt_1 dt_2}{\exp\left(\rho_g/2\right) \int_0^{\Delta} f_s(t) dt} \\
&= 1 + \frac{\exp\left(\rho_g/2\right) \int_0^{\Delta} \int_0^{\Delta} f_s(t_1) f_s(t_2) \left(\exp\left(K_g(t_1, t_2)\right) - 1\right) dt_1 dt_2}{\int_0^{\Delta} f_s(t) dt}.
\end{aligned}
$$

In the special case where the stimulus component is non-fluctuating, i.e., $f_s(t) = \bar{f}$, these expressions simplify to:

$$
\begin{aligned}
\mathbb{E}\left[y_\Delta \mid s\right] &= \exp\left(\rho_g/2\right) \int_0^{\Delta} f_s(t) dt \\
&= \exp\left(\rho_g/2\right) \bar{f} \Delta \\
&= \Delta \bar{\lambda},
\end{aligned}
$$

$$
\begin{aligned}
\operatorname{var}[y_\Delta \mid s] &= \exp\left(\rho_g/2\right) \int_0^{\Delta} f_s(t) dt + \exp\left(\rho_g\right) \iint f_s(t_1) f_s(t_2) \left(\exp\left(K_g(t_1, t_2)\right) - 1\right) dt_1 dt_2 \\
&= \exp\left(\rho_g/2\right) \bar{f} \Delta + \exp\left(\rho_g\right) \bar{f}^2 \left(\iint \left(\exp\left(K_g(t_1, t_2)\right)\right) dt_1 dt_2 - \Delta^2\right) \\
&= \Delta \bar{\lambda} + \left(\Delta \bar{\lambda}\right)^2 \times \left(\frac{1}{\Delta^2} \iint \left(\exp\left(K_g(t_1, t_2)\right)\right) dt_1 dt_2 - 1\right) \\
&= \Delta \bar{\lambda} + \left(\Delta \bar{\lambda}\right)^2 \left(\kappa(\Delta) - 1\right),
\end{aligned}
$$

and:

$$
FF(\Delta) = 1 + \Delta \bar{\lambda} (\kappa(\Delta) - 1),
$$

where we define $\bar{\lambda} = \bar{f} \exp(\rho_g/2)$ and $\kappa(\Delta) = \frac{1}{\Delta^2} \int_0^{\Delta} \int_0^{\Delta} e^{K_g(t_1, t_2)} \, dt_1 \, dt_2$.

#### 3 Details about the Goris model variants

In this appendix, we provide technical details for two extensions of the modulated Poisson framework introduced by Goris et al. [5], adapted for spike-train data with multiple time bins per trial. Specifically, we derive the likelihoods, means, and variances for the Goris-indep model, which assumes time-bin–independent modulatory gains, and the Goris-const model, which assumes a constant gain across time bins within a trial. These derivations clarify the statistical properties of each model and describe how we compute log-likelihoods for model comparison. All Goris-style baselines reported in the main model comparisons use a smooth stimulus-drive estimate obtained from the Poisson-GP model, so that comparisons among these baselines focus on the assumed temporal structure of the gain process. We also evaluate versions without this smoothness prior, in which the stimulus-drive parameters are estimated directly under the corresponding Goris-style likelihood. Details for both versions are provided below.

##### 3.1 Goris-indep model

Here we provide further details on the Goris-independent gain model, including the derivation of the likelihood, as well as the mean and variance under this model. The Goris-indep model is an extension of the modulated Poisson model in Goris et al. [5], adapted for spike train data with multiple time bins per trial. Given spiking observations of the $s^{\text{th}}$ stimulus presentation across $t = 1, \ldots, T$ time bins, we assume that the modulatory gain process is independent across time bins, with shared variance $\sigma^2_{g_s}$, i.e.,

$$\begin{aligned} y_{t,s,n} &\sim \mathrm{Poisson}(\lambda_{t,s,n}), \\ \lambda_{t,s,n} &= g_{t,s,n} f_{t,s} dt, \end{aligned}$$

where $g_{t,s,n} \sim \mathrm{Gamma}(r_s, s_s)$, with shape parameter $r_s = \frac{1}{\sigma^2_{g_s}}$ and scale parameter $s_s = \sigma^2_{g_s}$. This implies $\lambda_{t,s,n} \sim \mathrm{Gamma}\left(r_s, s_{t,s}\right)$, where $s_{t,s} = \sigma^2_{g_s} f_{t,s} dt$.

###### 3.1.1 Likelihood derivation

Following the results in [5], marginalizing over the gain variable yields independent Negative Binomial distributions:

$$\begin{aligned} & p(y_{1:T,s,1:N} | r_s, s_s) \\ &= \int_0^\infty p(y_{1:T,s,1:N}, \lambda_{1:T,s,1:N} | r_s, s_s)) d\lambda_{1:T,s,1:N} \qquad (5) \\ &= \int_0^\infty \prod_{t,n=1}^{T,N} \mathrm{Poisson}\left(y_{t,s,n} | \lambda_{t,s,n}\right) \times \mathrm{Gamma}\left(\lambda_{t,s,n} | r_s, s_{t,s}\right) d\lambda_{t,s,n} \\ &= \prod_{t,n=1}^{T,N} \int_0^\infty \exp\left(-\lambda_{t,s,n}\right) \frac{\lambda_{t,s,n}^{y_{t,s,n}}}{y_{t,s,n}!} \times \lambda_{t,s,n}^{r_s - 1} \frac{\exp\left(-\frac{\lambda_{t,s,n}}{s_{t,s}}\right)}{s_{t,s}^{r_s} \Gamma(r_s)} d\lambda_{t,s,n} \\ &= \prod_{t,n=1}^{T,N} \frac{1}{s_{t,s}^{r_s}} \frac{1}{y_{t,s,n}! \Gamma(r_s)} \int_0^\infty \lambda_{t,s,n}^{r_s + y_{t,s,n} - 1} \exp\left(-\lambda_{t,s,n} \left(1 + \frac{1}{s_{t,s}}\right)\right) d\lambda_{t,s,n} \end{aligned}$$

$$= \prod_{t,n=1}^{T,N} \frac{1}{s_{t,s}^{r_s}} \frac{1}{y_{t,s,n}!\Gamma(r_s)} \Gamma(r_s + y_{t,s,n}) \left( \frac{s_{t,s}}{1+s_{t,s}} \right)^{y_{t,s,n}+r_s} \int_0^\infty \text{Gamma}\left( \lambda_{t,s,n} \middle| r_s + y_{t,s,n}, \frac{s_{t,s}}{1+s_{t,s}} \right) d\lambda_{t,s,n}$$
$$= \prod_{t,n=1}^{T,N} \frac{\Gamma(r_s + y_{t,s,n})}{y_{t,s,n}!\Gamma(r_s)} \left( \frac{1}{1+s_{t,s}} \right)^{r_s} \left( \frac{s_{t,s}}{1+s_{t,s}} \right)^{y_{t,s,n}}$$
$$= \prod_{t,n=1}^{T,N} \frac{(r_s + y_{t,s,n} - 1)!}{y_{t,s,n}!(r_s - 1)!} \left( \frac{1}{1+s_{t,s}} \right)^{r_s} \left( \frac{s_{t,s}}{1+s_{t,s}} \right)^{y_{t,s,n}}$$
$$= \prod_{t,n=1}^{T,N} \text{NegativeBinomial}\left( y_{t,s,n} \middle| r_s, \frac{1}{1+s_{t,s}} \right).$$

For the Goris-indep model used in the main model comparisons, we set the stimulus component to the smooth firing-rate estimate obtained from the Poisson-GP model. Specifically, we set:

$$s_{t,s} = \sigma_{g_s}^2 f_{t,s} dt = f_{t,s} dt / r_s = \hat{\lambda}_{t,s} / r_s,$$

where $\hat{\lambda}_{t,s}$ denotes the Poisson-GP estimate of the stimulus-dependent firing rate. We then search for the value of $r_s$ that minimizes the negative log-likelihood of the spiking observations $y_{1:T,s,1:N}$ for each stimulus condition. For the comparison without a smoothness prior on the stimulus drive, we instead estimate both $r_s$ and $s_{1:T,s}$ directly by minimizing the negative log-likelihood under the above Negative Binomial mixture model, as in Goris et al. [5].

##### 3.1.2 Mean-variance derivations

Based on the above derivation, we see that:

$$p(y_{t,s,n} | r_s, s_{t,s}) \sim \text{NegativeBinomial}\left( r_s, \frac{1}{1+s_{t,s}} \right),$$

and $y_{1:t,s,n}$ are independent across $n = 1, \ldots, N$ when conditioned on $r_s$ and $s_{t,s}$. Thus, from the mean and variance of the Negative Binomial distribution, we have:

$$\begin{aligned} \text{mean}\left[ y_{t,s,1:N} | f_{t,s}, dt \right] &= \mathbb{E}\left[ y_{t,s,n} | f_{t,s}, dt \right] = r_s s_{t,s} = f_{t,s} dt, \\ \text{var}\left[ y_{t,s,1:N} | f_{t,s}, dt \right] &= r_s s_{t,s}(1 + s_{t,s}) = f_{t,s} dt \left( 1 + \sigma_{g_s}^2 f_{t,s} dt \right). \end{aligned}$$

Next, we extend this to derive the mean and variance of spike counts over a larger time window $\Delta = R\,dt$, by summing across the relevant bins $t_1, \ldots, t_R$:

$$\begin{aligned} \text{mean}\left( y_{\Delta,s,1:N} | \mathbf{f_s}, dt \right) = \mathbb{E}\left[ y_{\Delta,s,n} | \mathbf{f_s}, dt \right] &= \mathbb{E}\left[ \sum_{r'=1}^{R} y_{t_{r'},s,n} \middle| \mathbf{f_s}, dt \right] \\ &= \sum_{r'=1}^{R} \mathbb{E}\left[ y_{t_{r'},s,n} | \mathbf{f_s}, dt \right] \\ &= \sum_{r'=1}^{R} r_s s_{t_{r'},s} \end{aligned}$$

$$= \sum_{r'=1}^{R} f_{t_{r'},s} dt,$$

and:

$$\begin{aligned}
\text{var}\left[y_{\Delta,s,1:N} | \mathbf{f_s}, dt\right] &= \text{var}\left[\sum_{r'=1}^{R} y_{t_{r'},s,n} \middle| \mathbf{f_s}, dt\right] \\
&= \sum_{r'=1}^{R} \text{var}\left[y_{t_{r'},s,n} \middle| \mathbf{f_s}, dt\right] \\
&= \sum_{r'=1}^{R} r_s s_{t_{r'},s} \left(1 + s_{t_{r'},s}\right) \\
&= \sum_{r'=1}^{R} f_{t_{r'},s} dt \left(1 + \sigma_{g_s}^2 f_{t_{r'},s} dt\right).
\end{aligned}$$

##### 3.2 Goris-const model

In this section, we provide further details on the Goris-constant gain model, including the derivation of the mean and variance, as well as the log-likelihood under its modeling assumptions. The Goris-const model is the second variant we consider for extending the Goris model in [5] to multiple time bins. Here, we assume that the modulatory gain process is constant across time bins:

$$\begin{aligned}
y_{t,s,n} &\sim \text{Poisson}(\lambda_{t,s,n}), \\
\lambda_{t,s,n} &= g_{s,n} f_{t,s} dt,
\end{aligned}$$

where $g_{s,n} \sim \text{Gamma}(r_s, s_s)$, with shape parameter $r_s = \frac{1}{\sigma_{g_s}^2}$ and scale parameter $s_s = \sigma_{g_s}^2$. Furthermore, we assume that the total spike count across all time bins in a trial, $Y_{s,n} = \sum_t y_{t,s,n}$, shares the same gain variance $\sigma_{g_s}^2$, i.e.,

$$Y_{s,n} \sim \text{Poisson}(\tilde{g}_{s,n} f_s T), \tag{6}$$

with $\tilde{g}_{s,n} \sim \text{Gamma}(r_s, s_s)$. For the Goris-const model used in the main model comparisons, we estimate $\sigma_{g_s}^2$ by fitting the total spike counts $Y_{s,n}$ across training trials, as in Goris et al. [5], and set the stimulus encoding component $f_{t,s} dt = \hat{\lambda}_{t,s}$, where $\hat{\lambda}_{t,s}$ denotes the smooth firing-rate estimate obtained from the Poisson-GP model. For the comparison without a smoothness prior on the stimulus drive, we instead estimate $f_{t,s} dt$ using the same method as in the baseline Poisson model.

###### 3.2.1 Mean-variance derivations

Note that from the moments of the Gamma distribution:

$$\mathbb{E}\left[g_{s,n}\right] = 1, \quad \mathbb{E}\left[{g_{s,n}}^2\right] = \sigma_{g_s}^2 + 1. \tag{7}$$

Accordingly, we derive the first moment, second moment, and variance at bin size $dt$ as:

$$\text{mean}\left[y_{t,s,1:N} | f_{t,s}, dt\right] = \mathbb{E}\left[y_{t,s,n} | f_{t,s}, dt\right] = \mathbb{E}\left[\mathbb{E}\left[y_{t,s,n} | f_{t,s}, dt, g_{s,n}\right]\right] = \mathbb{E}\left[g_{s,n} f_{t,s} dt\right] = f_{t,s} dt \mathbb{E}\left[g_{s,n}\right]$$

$$
\begin{aligned}
&= f_{t,s}dt, \\
\mathbb{E}\left[y_{t,s,n}^2 | f_{t,s}, dt\right] &= \mathbb{E}\left[\mathbb{E}\left[y_{t,s,n}^2 | f_{t,s}, dt, g_{s,n}\right]\right] = \mathbb{E}\left[g_{s,n} f_{t,s} dt + g_{s,n}^2 f_{t,s}^2 dt^2\right] \\
&= f_{t,s}dt + f_{t,s}^2 dt^2(\sigma_{g_s}^2 + 1), \\
\mathbb{E}\left[y_{t_1,s,n} y_{t_2,s,n} | \mathbf{f_s}, dt\right] &= \mathbb{E}\left[\mathbb{E}\left[y_{t_1,s,n} y_{t_2,s,n} | \mathbf{f_s}, dt, g_{s,n}\right]\right] = \mathbb{E}\left[g_{s,n}^2 f_{t_1,s} f_{t_2,s} dt^2\right] = f_{t_1,s} f_{t_2,s} dt^2(\sigma_{G_s}^2 + 1), \\
\mathrm{var}\left[y_{t,s,1:N} | f_{t,s}, dt\right] &= \mathbb{E}\left[y_{t,s,n}^2 | f_{t,s}, dt\right] - \mathbb{E}\left[y_{t,s,n} | f_{t,s}, dt\right]^2 = f_{t,s}dt(f_{t,s}dt\sigma_{g_s}^2 + 1).
\end{aligned}
$$

We then extend these results to a larger bin size $\Delta = R\,dt$, by integrating over the differential statistics of the corresponding time bins $t_1, \cdots, t_R$ at bin-size $dt$:

$$
\begin{aligned}
\mathrm{mean}\left(y_{\Delta,s,1:N} | \mathbf{f_s}, dt\right) = \mathbb{E}\left[y_{\Delta,s,n} | \mathbf{f_s}, dt\right] &= \mathbb{E}\left[\sum_{r=1}^{R} y_{t_r,s,n} \middle| \mathbf{f_s}, dt\right] \\
&= \sum_{r=1}^{R} \mathbb{E}\left[y_{t_r,s,n} | \mathbf{f_s}, dt\right] \\
&= \sum_{r=1}^{R} f_{t_r,s} dt,
\end{aligned}
$$

$$
\begin{aligned}
\mathbb{E}\left[y_{\Delta,s,n}^2 | \mathbf{f_s}, dt\right] &= \mathbb{E}\left[\left(\sum_{r=1}^{R} y_{t_r,s,n}\right)^2 \middle| \mathbf{f_s}, dt\right] \\
&= \mathbb{E}\left[\sum_{r=1}^{R}\sum_{r'=1}^{R} y_{t_r,s,n} y_{t_{r'},s,n} \middle| \mathbf{f_s}, dt\right] \\
&= \sum_{r=1}^{R} \mathbb{E}\left[y_{t_r,s,n}^2 \middle| \mathbf{f_s}, dt\right] + \sum_{r \neq r'} \mathbb{E}\left[y_{t_r,s,n} y_{t_{r'},s,n} \middle| \mathbf{f_s}, dt\right] \\
&= \sum_{r=1}^{R} \left(f_{t_r,s}dt + f_{t_r,s}^2 dt^2(\sigma_{g_s}^2 + 1)\right) + \sum_{r \neq r'} \left(f_{t_r,s} f_{t_{r'},s} dt^2(\sigma_{g_s}^2 + 1)\right) \\
&= \sum_{r=1}^{R} f_{t_r,s}dt + \sum_{r=1}^{R}\sum_{r'=1}^{R} \left(f_{t_r,s} f_{t_{r'},s} dt^2(\sigma_{g_s}^2 + 1)\right),
\end{aligned}
$$

and therefore, the variance becomes:

$$
\begin{aligned}
\mathrm{var}\left[y_{\Delta,s,1:N} | \mathbf{f_s}, dt\right] &= \mathbb{E}\left[y_{\Delta,s,n}^2 | \mathbf{f_s}, dt\right] - \mathbb{E}\left[y_{\Delta,s,n} | \mathbf{f_s}, dt\right]^2 \\
&= \sum_{r=1}^{R} f_{t_r,s}dt + \sum_{r=1}^{R}\sum_{r'=1}^{R} \left(f_{t_r,s} f_{t_{r'},s} dt^2 \sigma_G^2\right) \\
&= \sum_{r=1}^{R} f_{t_r,s}dt \left(1 + \sigma_{g_s}^2 \sum_{r'=1}^{R} f_{t_{r'},s}dt\right).
\end{aligned}
$$

##### 3.2.2 Log-likelihood derivation

We calculate the test log-likelihood of this model using a Monte Carlo integration procedure [6]:

$$
\log\left(p\left(\mathbf{y} | \mathbf{f}\right)\right) = \sum_{k,s} \log\left(p\left(\mathbf{y}_{s,n} | \mathbf{f}_s\right)\right), \text{ where,} \tag{8}
$$

$$
\begin{aligned}
p\left(\mathbf{y}_{s,n}|\mathbf{f}_s\right) &= \int p\left(\mathbf{y}_{s,n}, g_{s,n}|\mathbf{f}_s\right) dg_{s,n} = \int p\left(\mathbf{y}_{s,n}|g_{s,n}, \mathbf{f}_s\right) p\left(g_{s,n}\right) dg_{s,n} \\
&= \mathbb{E}_{g_{s,n}}\left[p\left(\mathbf{y}_{s,n}|g_{s,n}, \mathbf{f}_s\right)\right] \\
&\approx \frac{1}{I}\sum_{i=1}^{I} p\left(\mathbf{y}_{s,n}|g_{s,n}^{(i)}, \mathbf{f}_s\right), g_{s,n}^{(i)} \sim \mathrm{Gamma}(r_s, s_s).
\end{aligned}
$$

Defining:

$$
\begin{aligned}
L_{s,n}^{(i)} &= \log\left(p\left(\mathbf{y}_{s,n}|g_{s,n}^{(i)}, \mathbf{f}_s\right)\right) \\
&= \log\left(\prod_{t=1}^{T} \frac{\exp\left(-g_{s,n}^{(i)} f_{t,s} dt\right)\left(g_{s,n}^{(i)} f_{t,s} dt\right)^{y_{t,s,n}}}{y_{t,s,n}!}\right) \\
&= \sum_{t=1}^{T}\left(-g_{s,n}^{(i)} f_{t,s} dt + y_{t,s,n}\log\left(g_{s,n}^{(i)} f_{t,s} dt\right) - \log\left(y_{t,s,n}!\right)\right),
\end{aligned}
$$

we get:

$$
\begin{aligned}
\log\left(p\left(\mathbf{y}_{s,n}|\mathbf{f}_s\right)\right) &\approx \log\left(\frac{1}{I}\sum_{i=1}^{I}\exp\left(L_{s,n}^{(i)}\right)\right) \\
&= \log\left(\sum_{i=1}^{I}\exp\left(L_{s,n}^{(i)} - L_{s,n}^{\mathsf{max}}\right)\right) + L_{s,n}^{\mathsf{max}} - \log\left(I\right),
\end{aligned}
$$

where $L_{s,n}^{\mathsf{max}} = \max_i L_{s,n}^{(i)}$. We set $I = 1000$ and use this formulation to derive the log-likelihood of the spiking observations in the $n^{\text{th}}$ test trial of the $s^{\text{th}}$ stimulus, and finally calculate the joint log-likelihood of all test data following Equation 8.

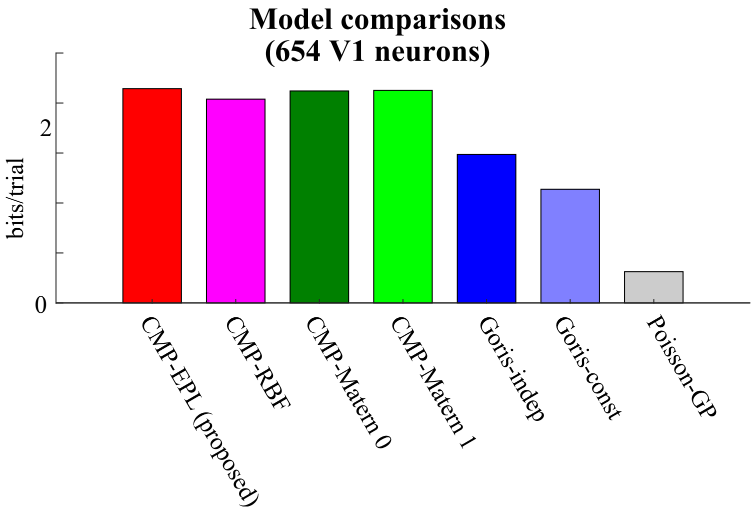


Figure S1: Comparison of the average test data log-likelihood improvement of each model relative to the baseline Poisson model, across 654 V1 neurons and all 72 grating orientations in recordings from the macaque primary visual cortex (data from [7]). From left to right: the proposed CMP model, the CMP-RBF model, the CMP-Matérn 0 model, the CMP-Matérn 1 model, the Goris-independent gain model, the Goris-constant gain model, and the Poisson-GP model. These results demonstrate that the CMP model systematically outperforms all other models in fitting held-out data.

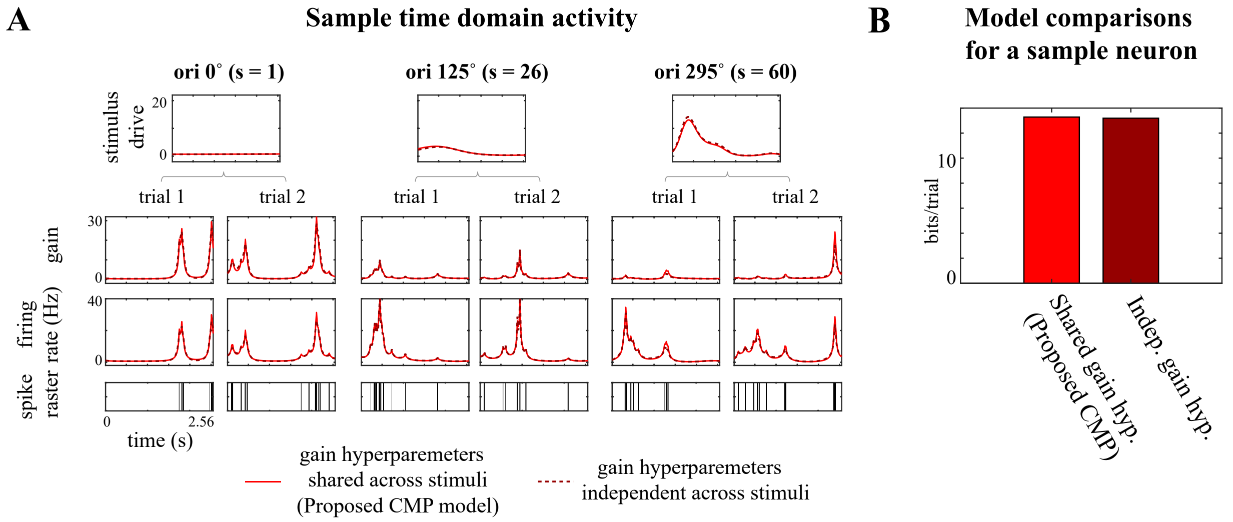


Figure S2: Comparison of the performance of the proposed CMP framework with an alternative model in which, instead of sharing a common set of gain hyperparameters $\rho_g, \ell_g, q$ across stimuli (as in the CMP framework), each stimulus condition has its own independent gain hyperparameters $\rho_g^{(s)}, \ell_g^{(s)}, q^{(s)}$. Results are shown for a sample neuron from the macaque primary visual cortex dataset (data from [7]). **(A)** Comparison of the inferred time-domain activity. From top to bottom: the inferred stimulus-driven components at three selected orientation settings ($0°$, $125°$, $295°$); inferred gain processes for two trials per orientation; inferred firing rates from the proposed CMP model (solid lines) and the alternative model with independent gain hyperparameters per stimulus (dashed lines); and the observed spike raster. **(B)** Comparison of the log-likelihood improvement relative to the baseline Poisson model for the proposed CMP model (left) versus the alternative independent-gain model (right). These results show that the performance of both models is very similar, corroborating that the proposed CMP modeling framework robustly captures trial-to-trial variability across different stimuli using a shared set of hyperparameters.

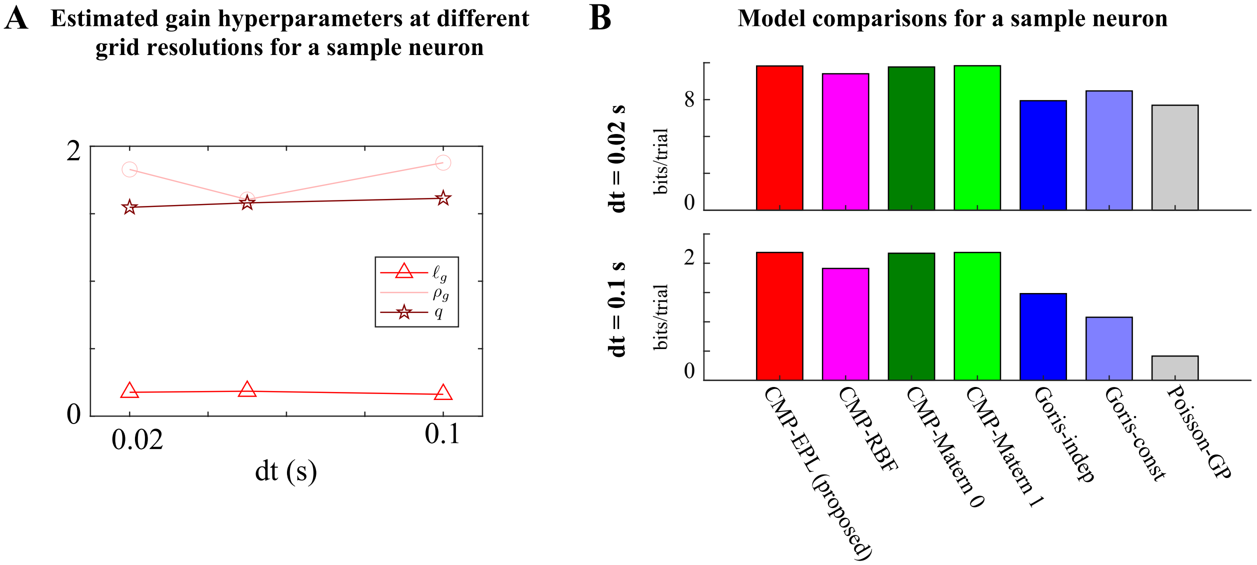


Figure S3: Comparison of CMP model inference at different grid sizes $dt$ for a sample neuron from the macaque primary visual cortex dataset (data from [7]). **(A)** Comparison of the inferred gain hyperparameters $\ell_g, \rho_g, q$ across three sufficiently fine grid resolutions: $dt = 0.02$ s, $dt = 0.05$ s, and $dt = 0.1$ s. These results indicate that the inference procedure is robust to moderate changes in grid size once the temporal discretization is fine enough to approximate the underlying continuous-time model. **(B)** Comparison of the test data log-likelihood gain relative to the baseline Poisson model at two different grid resolutions ($dt = 0.02$ s and $dt = 0.1$ s). From left to right: the proposed CMP model, CMP-RBF model, CMP-Matérn 0 model, CMP-Matérn 1 model, Goris-independent gain model, Goris-constant gain model, and the Poisson-GP model. These results show that the CMP consistently outperforms alternative models over this range.

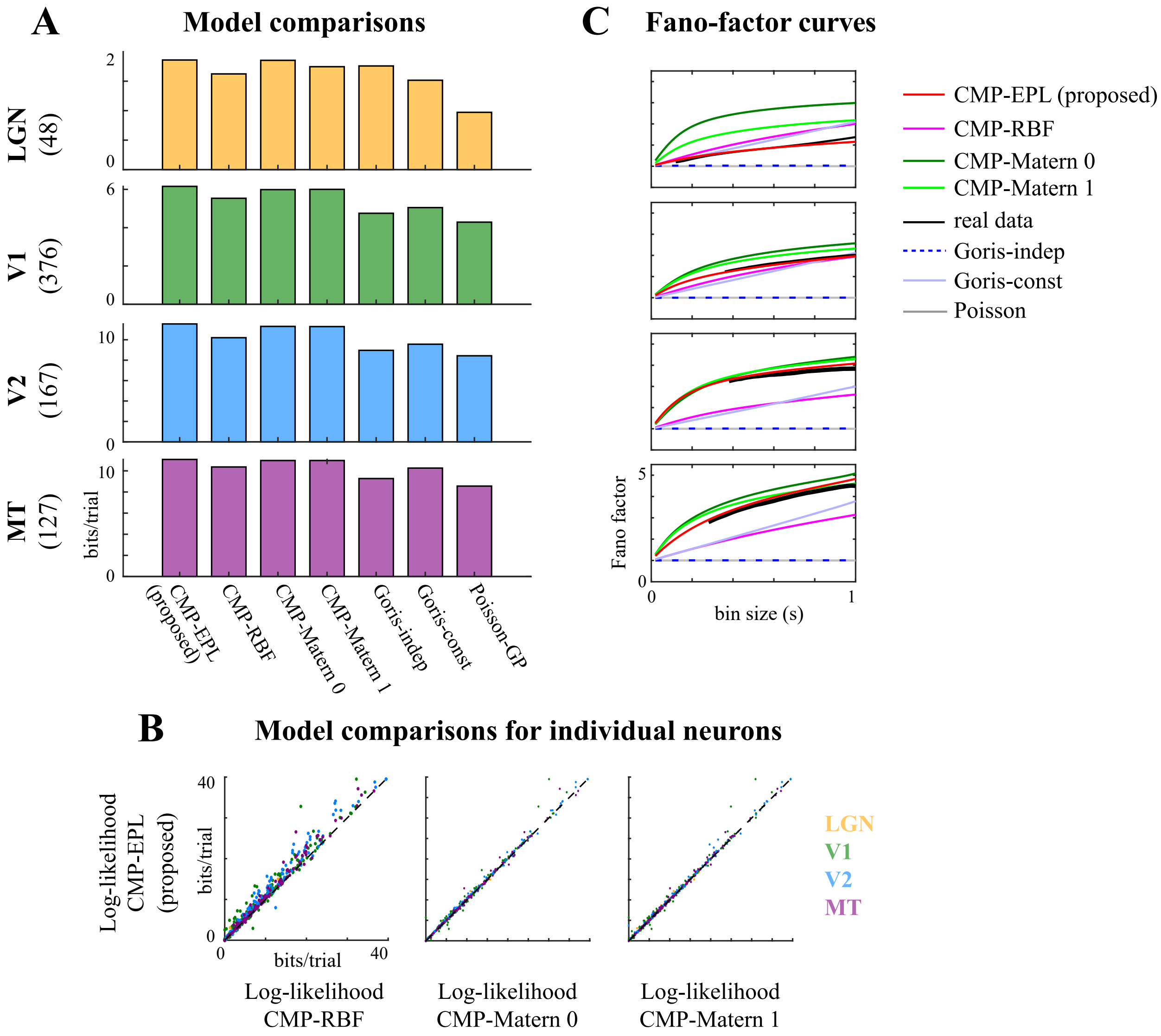


Figure S4: Performance comparisons of different models on neuronal population data recorded from various areas along the visual hierarchy: the lateral geniculate nucleus (LGN), V1, V2, and MT, in response to drifting sinusoidal gratings of the preferred size and speed, varying either in spatial frequency (12 spatial frequencies, ranging from 0 to 10 cycles/deg) or in drift direction (16 equally spaced directions) (data from [5]). **(A)** Log-likelihood improvement of each model relative to the baseline Poisson model, averaged across held-out data for each population. Rows from top to bottom: LGN, V1, V2, and MT. The number of neurons in each population is indicated in parentheses. From left to right: proposed CMP model, CMP-RBF model, CMP-Matérn 0 model, CMP-Matérn 1 model, Goris-independent gain model, Goris-constant gain model, and the Poisson-GP model. **(B)** Comparison of the log-likelihood of the proposed CMP model with the CMP-RBF model (left), the CMP-Matérn 0 model (middle), and the CMP-Matérn 1 model (right) for individual neurons. Colors indicate brain areas: LGN (yellow), V1 (green), V2 (blue), and MT (purple). **(C)** Comparison of the inferred Fano factor as a function of bin size, averaged across each population (rows from top to bottom: LGN, V1, V2, and MT). Curves show the real data (black), the proposed CMP model (red), CMP-RBF model (magenta), CMP-Matérn 0 model (dark green), CMP-Matérn 1 model (light green), Goris-independent gain model (dark blue dashed lines), Goris-constant gain model (light blue), and the baseline Poisson model (gray).

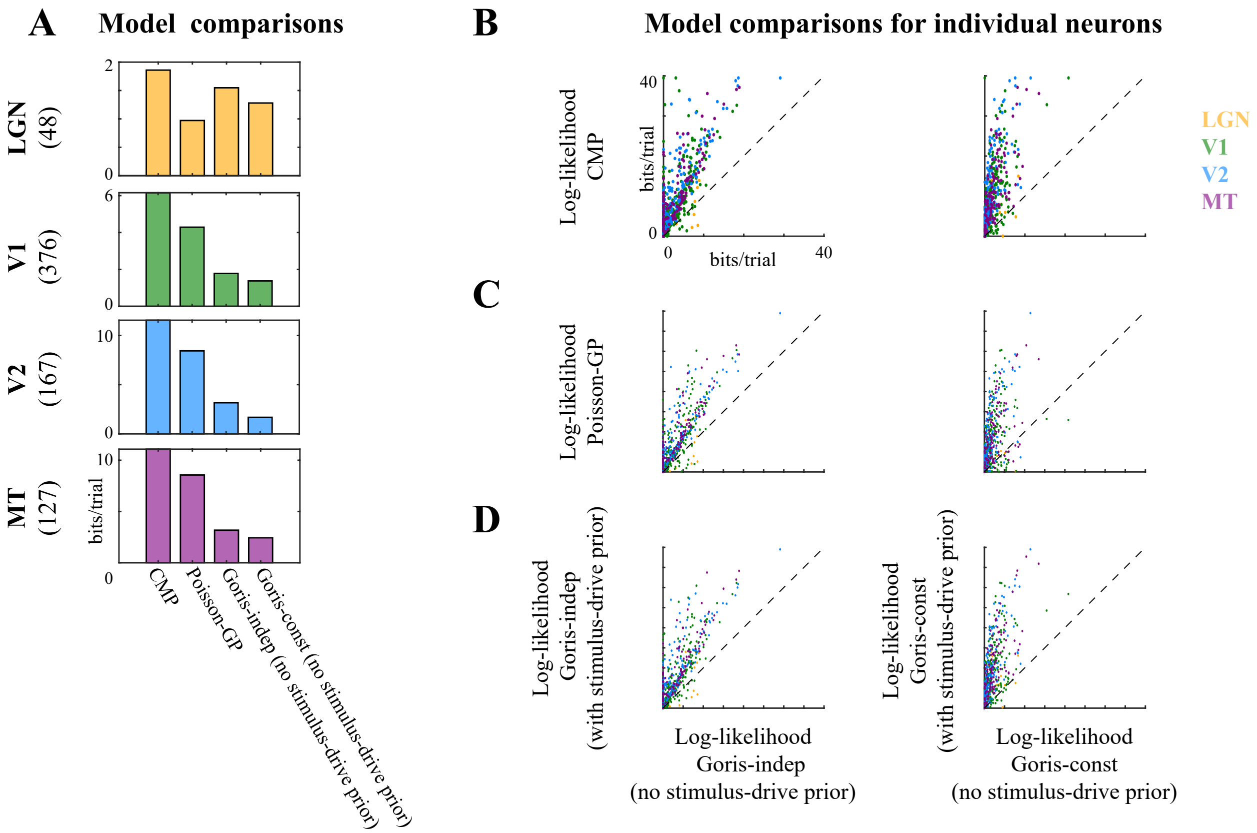


Figure S5: Comparison of the performance of the Goris model with and without a smoothness prior on the stimulus drive (data from [5]). **(A)** Log-likelihood improvement of each model relative to the baseline Poisson model, averaged across held-out data for each population. Rows from top to bottom: LGN, V1, V2, and MT, with the number of neurons in each population indicated in parentheses. From left to right: the proposed CMP model, the Poisson-GP model, the Goris-independent gain model without a GP prior on the stimulus drive, and the Goris-constant gain model without a GP prior on the stimulus drive. **(B)** Comparison of the log-likelihood of the proposed CMP model versus the Goris-independent gain model without a GP prior on the stimulus drive (left), and versus the Goris-constant gain model without a GP prior on the stimulus drive (right), for individual neurons. **(C)** Comparison of the log-likelihood of the Poisson-GP model versus the Goris-independent gain model without a GP prior (left), and versus the Goris-constant gain model without a GP prior (right), for individual neurons. **(D)** Comparison of the log-likelihood of the Goris-independent gain model with a GP prior on the stimulus drive versus without a GP prior (left), and the Goris-constant gain model with a GP prior versus without a GP prior (right), for individual neurons. Colors indicate the areas: LGN (yellow), V1 (green), V2 (blue), and MT (purple). These results show that including a smoothness prior on the stimulus drive generally improves the model's ability to fit held-out data, though the proposed CMP framework still achieves systematically higher performance across all areas. This highlights the importance of jointly modeling smooth stimulus tuning and structured gain fluctuations for accurately capturing neural variability.

### References


[1] Carl Edward Rasmussen and Christopher K. I. Williams. *Gaussian processes for machine learning.* Adaptive computation and machine learning. MIT Press, 2006. ISBN 026218253X.

[2] I. J. Schoenberg. Metric spaces and positive definite functions. *Transactions of the American Mathematical Society*, 44(3):522–536, 1938. ISSN 00029947, 10886850. URL http://www.jstor.org/stable/1989894.

[3] G.B. Folland. *Real Analysis: Modern Techniques and Their Applications*. Pure and Applied Mathematics: A Wiley Series of Texts, Monographs and Tracts. Wiley, 2013. ISBN 9781118626399. URL https://books.google.com/books?id=wI4fAwAAQBAJ.

[4] George Casella and Roger L. Berger. *Statistical Inference*. Duxbury Press, Pacific Grove, CA, 2nd edition, 2002.

[5] R L T Goris, J A Movshon, and E P Simoncelli. Partitioning neuronal variability. *Nature Neuroscience*, 17(6):858–865, 2014.

[6] Christian P Robert, George Casella, and George Casella. *Monte Carlo statistical methods*, volume 2. Springer, 1999.

[7] A B A Graf, A Kohn, M Jazayeri, and J A Movshon. Decoding the activity of neuronal populations in macaque primary visual cortex. *Nature Neuroscience*, 14(2):239–245, Feb 2011. doi: 10.1038/nn.2733. URL https://doi.org/10.1038/nn.2733.
